# Organisms use mode-switching to solve the explore-vs-exploit problem

**DOI:** 10.1101/2023.01.11.523566

**Authors:** Debojyoti Biswas, Andrew Lamperski, Yu Yang, Kathleen Hoffman, John Guckenheimer, Eric S. Fortune, Noah J. Cowan

## Abstract

The inescapable link between sensing and movement generates a conflict between producing costly movements for gathering information (“explore”) versus using previously acquired information to achieve a goal (“exploit”). Determining the optimal balance between explore and exploit is a computationally intractable problem, necessitating the use of heuristics. We looked to nature to measure and model the solutions used by organisms. Here we show that the electric fish *Eigenmannia virescens* uses a salience-dependent mode-switching strategy to solve the explore–exploit conflict during a refuge tracking task. The fish produced distinctive non-Gaussian (i.e., non-normal) distributions of movement velocities characterized by sharp peaks for slower, task-oriented “exploit” movements and broad shoulders for faster, “explore” movements. The measures of non-normality increased in relation to increased sensory salience. Data from ten phylogenetically diverse organisms, from amoebae to humans, revealed the same distinctive distribution of movement velocities that were also modulated in relation to sensory salience. We propose a state-uncertainty based mode-switching heuristic that (1) reproduces the distinctive velocity distribution, (2) rationalizes modulation by sensory salience, and (3) outperforms the classic persistent excitation approach while using less energy. This mode-switching heuristic provides insights to purposeful exploratory behaviors in organisms as well as a framework for more efficient state estimation and control of robots.

## Main

Organisms display complex patterns of movement that arise from the interplay between obtaining information (“explore”)^1–3^ and using current information (“exploit”).^4^ Exploratory movements to gain information, and exploitative movements to achieve the task at hand, are often mediated by the same motor systems. For example, the weakly electric glass knifefish (*Eigenmannia virescens*) produces both information-seeking exploratory movements^2, 3, 5^ and goal-driven exploitative movements to remain within a refuge^6, 7^ using the same ribbon-fin locomotor system.^8, 9^ Both of these types of movement occur in a single linear dimension, along the rostrocaudal axis. This behavior makes *E. virescens* an excellent model system with which to investigate the interplay between explore and exploit movements: within a fixed refuge, fish produce ancillary back and forth exploratory movements to sense the refuge,^2, 3, 5^ but these back and forth (explore) movements conflict with the corrective movements (exploit) required for station keeping.

Resolving this conflict between explore movements^5^ versus goal-directed exploit movements is a computationally intractable optimization problem.^10–12^ How do organisms resolve the explore–exploit conflict? A simple heuristic to solve this problem would be for an organism to perform goal-directed exploit movements while superimposing continuous small exploratory sensing movements—in other words to use a persistent excitation approach.^13^ Indeed, this heuristic has proven effective (if suboptimal) as an engineering approach to solve the explore–exploit problem of identifying states and parameters of a dynamical system during task execution.^13^ If organisms were to employ such a strategy, they would produce movement statistics that correspond to a single behavioral mode (e.g., a single component Gaussian distribution) that continuously superimposes explore and exploit behavior.

In contrast, we discovered that *E. virescens* does not use a persistent excitation strategy; instead, it exhibits a mode-switching strategy between fast (explore) and slow (exploit) behavioral modes. This mode switching is modulated by sensory salience (Fig. 1–2). To assess the generality of this mode-switching strategy we investigated ten additional tasks performed by ten species ranging from amoebae to humans,^14–22^ using five major sensing modalities–vision, audition, olfaction, tactile sensing and electrosensation (Fig. 4). Based on this extensive reanalysis we found that such mode switching—and its dependence on sensory salience—is found across diverse behaviors, taxa, and sensing modalities (Fig. 4).

**Fig. 1.**
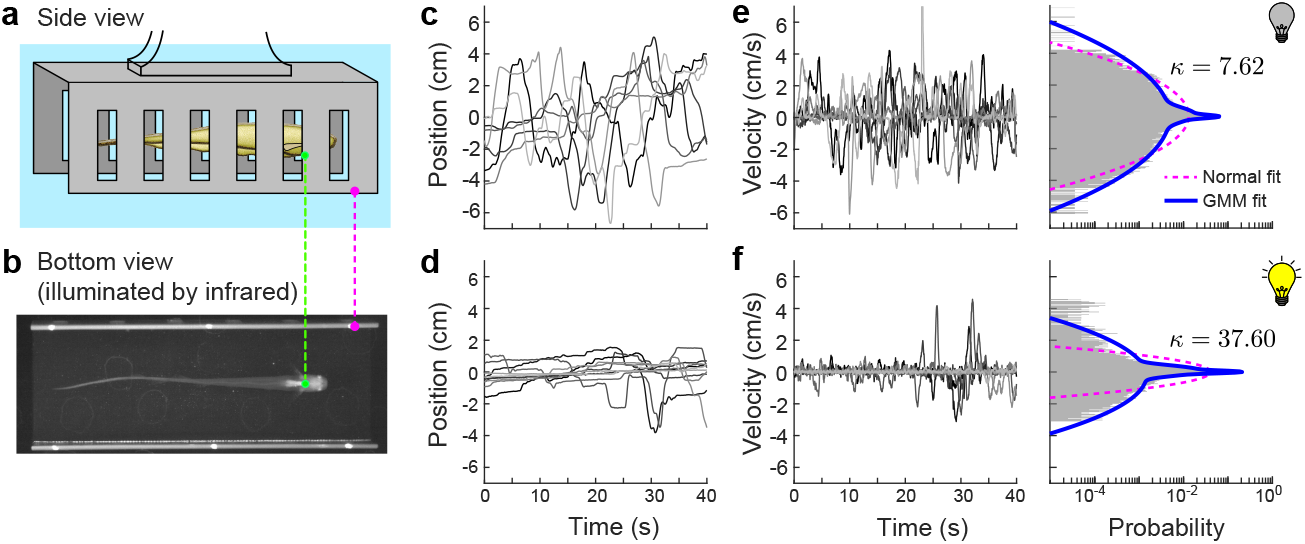
Velocity distributions are broad-shouldered. (a) Side view (schematic) and (b) bottom view (infrared image) of a fish inside a stationary refuge. The bright ventral patch on the fish was tracked (green dot) as the fish swam inside the refuge (magenta dot). (c,d) Position traces during (c) lights-off trials (*n* = 7), and (d) lights-on trials (*n* = 10) from a single representative fish. (e,f) Corresponding velocity traces during lights-off trials (left) and velocity histogram (right) over the same range of velocities for (e) lights-off, and (f) lights-on trials, with the kurtosis value *κ* indicated; note that the time and probability scales of the horizontal axes are shown below panel (f). A three-component Gaussian mixture model (GMM; blue solid curve) fit the data better than the normal fit (magenta dashed curve) as indicated by the lower Kullback-Leibler divergence and Bayesian information criterion values; see Extended Data Table 1 for statistical details. One lights-off trial with large positive velocity is truncated; see Extended Data Fig. 1a,b for full version.

**Fig. 2.**
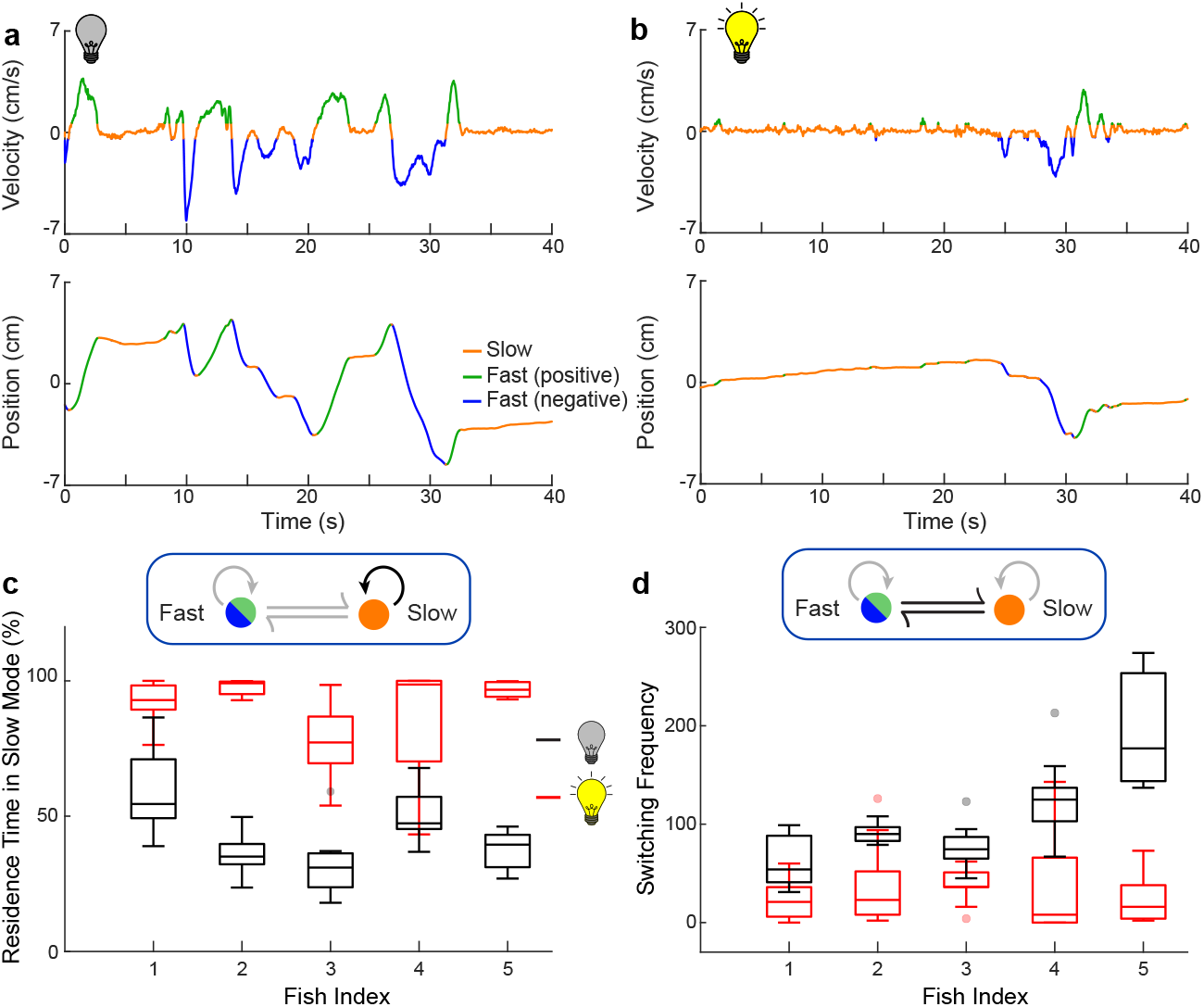
Bursts of faster movements are more common in lights-off trials than in lights-on trials. (a,b) Fish showed two distinct behavioral modes, slow movement and fast movement, as seen in the velocity (top) and position traces (bottom) of representative trials from the same fish under lights-off (a) and lights-on (b) conditions. (c) The residence time in the slow mode, computed as the percent of the trial duration (40 s), was significantly higher during lights-on trials than lights-off trials (one sided p-value *<* 0.002 for each fish). (d) The switching between the fast mode (positive and negative combined) and slow mode was significantly more frequent in lights-off trials (black) than in lights-on trials (red) (one-tailed p-value *<* 0.002 for each fish). For both (c,d), the p-values were calculated using the Mann-Whitney-Wilcoxon test.

Inspired by this widespread biological solution to the explore–exploit problem, we propose an engineering heuristic for selecting modes based on state uncertainty (Fig. 5) that captures the key structural features of mode switching found across organismal models. Furthermore, we show that this mode-switching heuristic can achieve better task-level performance, and do so with less control effort, than the conventional persistent excitation strategy.

### *E. virescens* exhibited fast and slow behavioral modes

We examined the behavior of individual *E. virescens* as they performed untrained station keeping within a fixed refuge. Station keeping requires only small corrective movements; therefore any significant movements by the fish are attributed to information-seeking, exploratory movement.^2, 3, 5^ Previous work^23^ has demonstrated that *E. virescens* use both vision and electrosense for station-keeping. Hence varying light level is an experimental mechanism to examine the effect of visual salience on the selection between explore and exploit movements.

We measured the movements of five individual fish in 40 s duration station-keeping trials, in two lighting conditions: lights “on” trials had bright illumination (*∼*80 lx), and lights “off” trials had low illumination (*∼*0.3 lx). We conducted between 7 and 10 trials per condition per fish. We discarded trials in which the fish changed its swimming direction or exited the refuge. Consistent with prior studies,^2, 5^ fish moved significantly more in lights-off conditions than in lights-on conditions (Extended Data Fig. 1). However, these previous analyses^2, 5^ focused on tracking performance using analytical methods including Fourier analysis and root-mean-square metrics that masked the temporal structure of active sensing movements that we seek to understand in this paper.

We found the patterns of fish swimming velocities were consistent with a mode-switching strategy. The distribution of velocities (*v*) featured a sharp peak around *v* = 0 with “broad” shoulders for faster movements (Fig. 1e,f, right). These empirical distributions differed from a Gaussian distribution in two ways: (i) the distinct central peak, and (ii) the broad shoulders corresponding to the faster movements. The central peak (near zero velocity) represents slow movement, and the broad shoulders represent faster movement. These behavioral modes are associated with exploit and explore respectively, as discussed in greater detail below.

The two behavioral modes were significantly better approximated by three-component Gaussian mixture models (GMMs) than by single component models (Fig. 1e,f, right). This was shown by three measures, namely Kullback-Leibler (K-L) divergence, Bayesian information criterion (BIC), and closeness of quantile-quantile (Q-Q) plots to the reference line (Extended Data Table 1, Extended Data Fig. 1). The three component GMMs generally comprised a sharp central Gaussian peak, capturing slow, task-oriented station-keeping movements, and two Gaussian “shoulders”, capturing faster, positive (forward) and negative (backward) exploratory movements. We found that only modest improvements in the fit of the GMMs occurred when using more than three components (Extended Data Fig. 1f).

The fast mode movements increased in frequency in lights-off trials, increasing the relative prominence of the “shoulders”. For example, Fig. 1e,f show representative data from one fish in which there were 48 fast movements with lights off (Fig. 1e, left) but only 13 fast movements with lights on (Fig. 1f, left). Interestingly, the overall higher proportion of fast velocities in lights-off trials leads to a surprising result, namely higher kurtosis values for lights-on versus lights-off trials (Extended Data Fig. 1e). In other words, the increase in frequency of fast motions in the dark leads to a decrease in the relative prominence of the central, task-oriented velocity peak at *v* = 0, so that the overall distribution is closer to Gaussian, and the kurtosis trends toward 3.

We found that the trend towards a Gaussian distribution of movement velocities in lights-off trials (reduced sensory salience) to be surprising because the exploratory movements for actively sensing the environment are associated with a nonlinear requirement^24, 25^ to make movements that are potentially in conflict with task goals. Therefore, our initial hypothesis– that this nonlinearity would produce *increased* deviation from a Gaussian velocity distribution as sensory salience was *reduced* –was not supported. Our initial intuition failed because we did not appreciate that decreases in sensory salience drives the selection of explore behavior, and that behavior itself is approximately Gaussian, ultimately reducing the relative prominence of the task-oriented central peak.

Interestingly, reanalysis of data from a previous study of exploratory movements in a similar refuge tracking paradigm in *E. virescens* show the same relationship between sensory salience and changes in velocity profiles, but for modulations of a different sensory modality, namely electrosensation.^3^ In these previous experiments, artificially generated electrical signals were used to diminish the salience of electrosensory information as the electric fish performed the refuge tracking. Our reanalysis of these published data (see Supplementary Material and Methods for details) showed that fish exhibited the distinctive non-Gaussian distribution of velocities. Moreover, the velocity distributions were modulated in relation to electrosensory salience: lower kurtosis values (corresponding to more normal distributions) occurred in experimental trials with added artificial electrosensory “jamming” signals (Extended Data Fig. 2a–f).

### Sensory salience drives explore–exploit mode switching

How do changes in sensory salience drive changes in mode switching? To investigate this question, we segregated the velocity trajectories into “S”, a slow velocity mode (exploit) comprising task-oriented, station-keeping movements, and “F”, a fast velocity mode (explore) comprising large positive (forward) and negative (backward) movement velocities (Fig. 2a,b, Extended Data Fig. 3) (see Methods for different clustering algorithms used).

Fish produced slow and fast velocity modes of movements in both lights-on and lights-off trials. We computed the residence time in each behavioral mode as a proportion of the total time spent in that mode compared to the trial duration of 40 s (note that the residence time in slow and fast modes adds up to unity). The residence time *τ_s_* in the slow exploit mode was significantly higher (*>* 1.7 times) in lights-on trials than the lights-off trials (Fig. 2c). In contrast, the residence time in the fast explore mode (1 *− τ_s_*) was higher in light-off than in lights-on trials.

Fish switched between slow and fast modes more frequently in lights-off trials than in lights-on trials (Fig. 2d). From the computation of the transition rates between slow (S) and fast (F) modes as a two-state Markov process, we found that the transition rate S *→* F was significantly lower in lights-on versus lights-off trials, i.e., the slow (exploit) state was visited more frequently in the lights-on trials compared to lights-off trials (Extended Data Fig. 3b). This salience-dependent modulation of switching frequency was the key mechanism by which movement velocity distributions trended toward a Gaussian distribution as a function of decreased sensory salience.

### Mode-switching is found across taxa, behaviors, and sensory modalities

Is this mode-switching strategy solution for the explore versus exploit problem found in other species, in other categories of behavioral tasks, and in control systems that rely on other sensing modalities?

To answer this question we analyzed published data for an additional ten species, representing a wide phylogenetic range of taxa, from single-celled organisms to humans, involving categorically different tasks and sensorimotor regimes.^3, 14–20, 22^ These taxonomically diverse species were selected to encompass a wide range of behaviors that rely on a broad range of sensory systems (Fig. 3). For every example we examined, we found the same distinctive non-normal distribution of velocities, with a peak at low velocity movements and broad shoulders for higher velocity movements (Fig. 4).

**Fig. 3.**
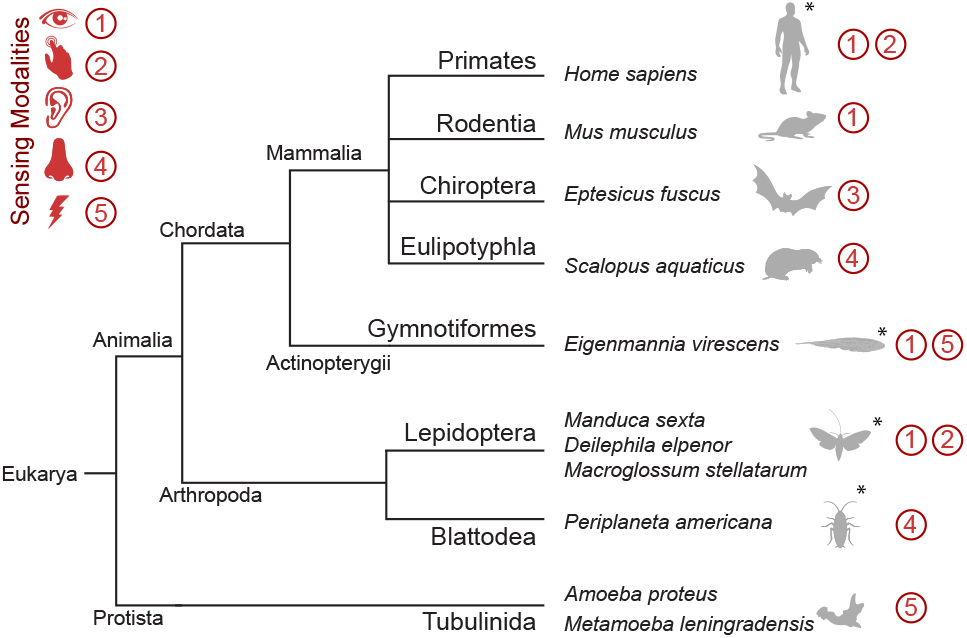
Diversity of organisms and sensing modalities used for analysing mode-switching strategies. The major sensing modalities (vision, tactile sensing, audition, olfaction, and eletrosensation) used for task execution are listed next to the organisms (four species of mammals, a species of ray-finned fish, four species of insects, and two species of amoebae). The effect of sensory salience is studied for organisms marked with asterisks. Behaviors and movement statistics of the organisms are shown in greater detail in Fig. 4.

For example, postural sway movements in humans are thought to prevent the fading of postural state information during balance.^26^ Our reanalysis of quiescent stance data^17^ revealed evidence of mode switching (Fig. 4a) that is remarkably similar to our findings in electric fish. In the quiescent stance task, human subjects used visual and tactile feedback to maintain an upright posture. The distribution of sway velocities revealed a distinct peak at low velocities corresponding to the task goal, and broad shoulders for higher velocities produced by the exploratory movements; the velocity statistics were better captured by a GMM than a normal distribution. Furthermore, the velocity distribution showed the same surprising relation to changes in salience, becoming more Gaussian as well as increase in the switching frequency when sensory salience was decreased (Extended Data Fig. 4a–d), as seen in the electric fish *E. virescens*.

**Fig. 4.**
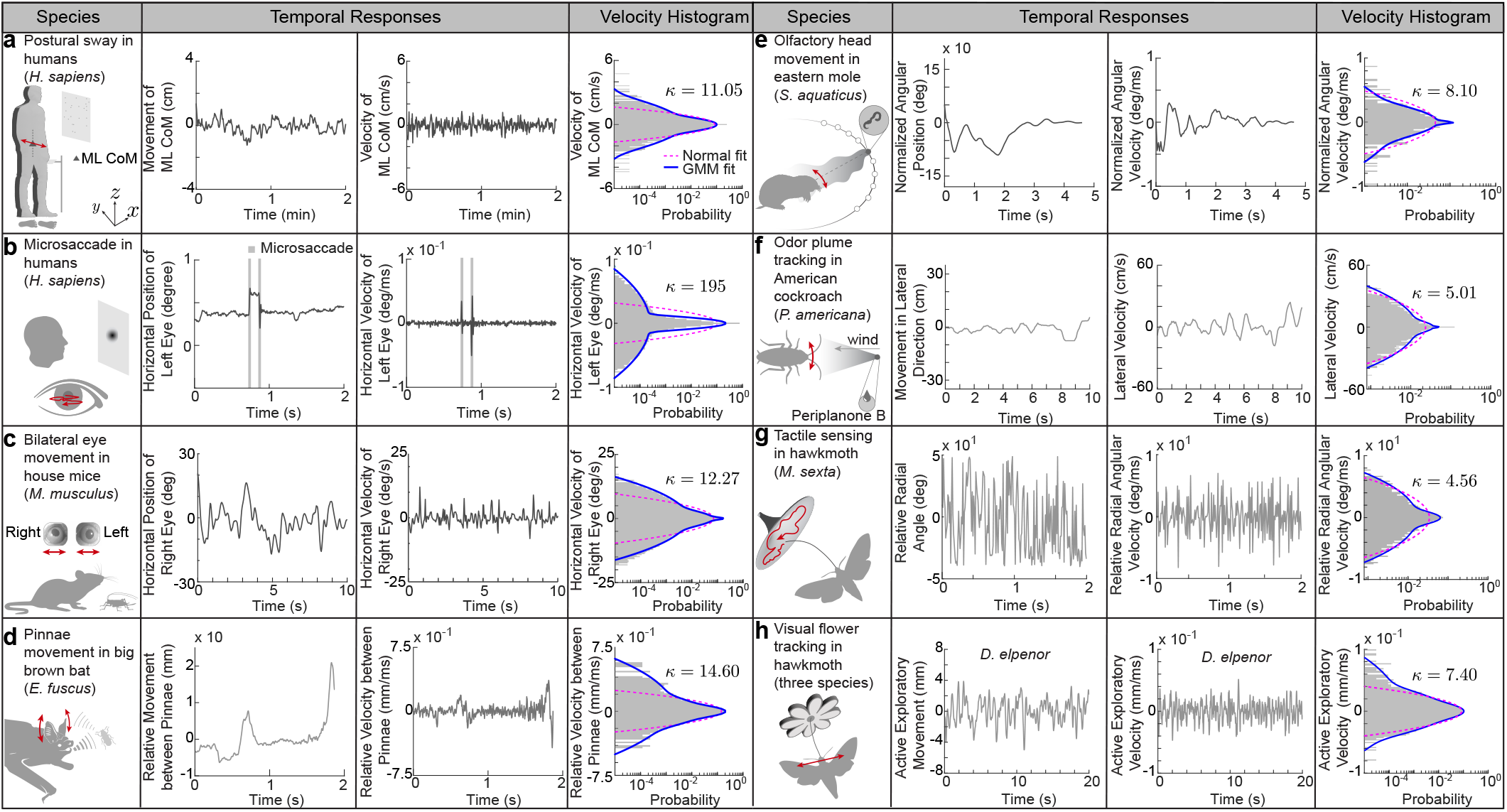
Broad-shouldered velocity distribution is found across taxa, behaviors, and sensing modalities. Reanalysis of data from eight prior studies reveals a convergent statistical structure of movements across a range of organisms and behaviors: (a) postural sway in humans (*Homo sapiens*) during maintenance of quiet upright stance,^17^ (b) microsaccades in humans (*Homo sapiens*) during fixated gaze,^21^ (c) bilateral eye movements in mice (*Mus musculus*) during prey (cricket) capture,^15^ (d) pinnae movements in big brown bats (*Eptesicus fuscus*) while echolocating prey (mealworm),^16^ (e) olfactory driven head movements in eastern moles (*Scalopus aquaticus*) in response to food (earthworms),^14^ (f) odor plume tracking in American cockroaches (*Periplaneta americana*) in response to sex pheromone (Periplanone B),^18^ (g) tactile sensing by Carolina sphinx hawkmoth (*Manduca sexta*) while searching for a flower nectary,^20^ and (h) visual tracking of swaying flower by hawkmoths^19^ (three species; only data from elephant hawkmoth, *Deilephila elpenor* is shown, for the remainder see Extended Data Fig. 5g–p).The second column shows representative temporal traces of the active exploratory movements, the third column shows the respective velocity traces. The fourth column presents velocity histograms showing that, unlike the normal distribution (magenta dashed curve), the three-component Gaussian mixture model (GMM; blue solid curve) captures the broad-shouldered nature of the velocity data across species, behaviors, and sensing modalities. See Extended Data Table 2 for statistical details. Photo credit: Mice eye image courtesy of authors in Michaiel *et al.*,^15^ published under the terms of the Creative Commons Attribution License.

Mode switching was also observed in invertebrate species. For example, the Carolina sphinx hawkmoth (*Manduca sexta*) uses somatosensory feedback from their proboscis to detect the curvature of flowers when searching for nectaries at dawn and dusk.^20^ In this search behavior, which has dynamics that are qualitatively similar to vibrissal sensing in rats,^27^ the moth sweeps its proboscis across the surface of the flower using a combination of slow- and high-velocity movements. Our analysis of the distribution of the rate of change of radial orientation angle (angle between the proboscis tip trajectory and the radial axis of the flower), before the insertion of the proboscis tip into the nectary shows the characteristic sharp peak with broad-shoulders (Fig. 4g) that is captured by a GMM. Experimental changes in the shape of artificial flowers that degrade the salience of the curvature of the flower surface^20^ resulted in a decrease of the kurtosis value of the proboscis angular velocity distribution (Extended Data Fig. 5a–f), similar to how both *E. virescens* and humans responded to changes in sensory salience.

The fact that this salience-based, mode switching strategy was found in two distantly related classes (mammalia and insecta) performing very different behaviors, using different sensorimotor systems, suggests that the strategy emerged as a convergent solution to the explore versus exploit problem. We found additional evidence of convergence of this solution in reanalysis of eight additional datasets: visual saccades in humans (Fig. 4b, Extended Data Fig. 7)^21^ and in house mice *Mus musculus* (Fig. 3c),^15^ movements of the pinnae of echolocating big brown bats *Eptesicus fuscus* (Fig. 4d),^16^ olfaction in Eastern moles *Scalopus aquaticus* (Fig. 4e)^14^ and American cockroaches *Periplaneta americana* (Fig. 4f, Extended Data Fig. 4e–f),^18^ and visual tracking of a swaying flower in three species of hawkmoths (*Manduca sexta, Deilephila elpenor, Macroglossum stellatarum*) (Fig. 4h, Extended Data Fig. 5 g–p).^19^ The discovery of a similar, parsimonious velocity distributions across taxa, behavior, and sensing modalities, with consistent dependency on sensory salience, was surprising.

Intriguingly, our analysis of the dynamics of transverse exploration by pseudopods of amoebae^22^ (*Amoeba proteus* and *Metamoeba leningradensis*) reveals similar GMM velocity distributions in response to an electric field (Extended Data Fig. 6). Although our modeling approach (see next section) includes inertial dynamics, that cannot be directly applied to movement of organisms in the low Reynolds number regimes occupied by single-celled and other microscopic organisms, these observations are consistent with a mode-switching strategy for the control of movement in these amoebae.

The examples described above include a broad phylogenetic array of organisms that perform a variety of behavioral tasks using different control and morphophysiological systems. Just as these behavioral systems evolved within each of the lineages represented in our reanalyses, we suggest that mode switching likely evolved independently in each lineage as well. In other words, the similarities we found across taxa are the result of convergent evolution towards a common solution—mode switching—for the explore versus exploit problem.

### Heuristic Model of the Explore–Exploit Mode-Switching Strategy

Why might animals use mode switching, rather than the simpler heuristic of applying continual, low-amplitude exploratory inputs used by control engineers^13^? To address this question, we propose a parsimonious heuristic model that comprises a nonlinear motion-dependent sensor, a linear musculoskeletal plant, a state estimator, and a mode-switching controller (Fig. 5a). For the musculoskeletal plant, we assumed a simplified second-order Newtonian model:^9, 24^

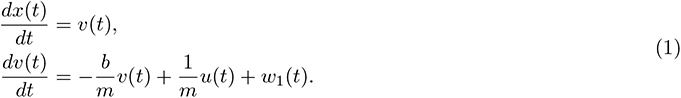

Here *x*(*t*) is dimensionless position, *v*(*t*) is dimensionless velocity, *u*(*t*) is the controller input, and *w*_1_(*t*) is process noise. The process noise includes noise due to physical disturbances^28, 29^ as well as motor noise.^30^ The system parameters *m* and *b* represent unitless mass and viscous damping, respectively.

**Fig. 5.**
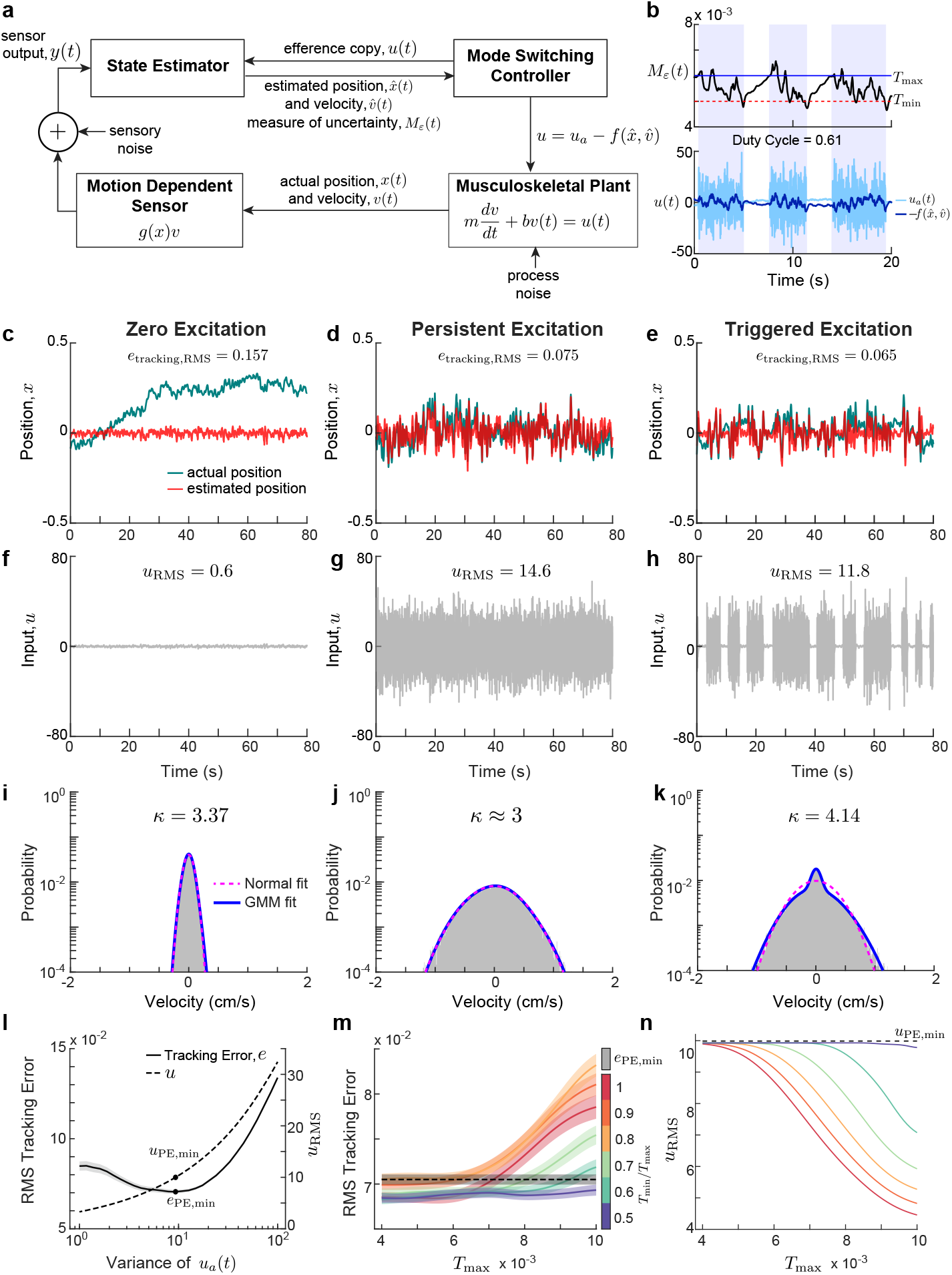
Template model illustrating three different exploration strategies. (a) Schematic of the Triggered-Excitation (mode-switching) strategy. The musculoskeletal plant for animal locomotion has two states— position (*x*) and velocity (*v*). The state estimator has access to noisy measurements from a nonlinear adaptive sensor (*g*(*x*)*v*). The state estimator (extended Kalman filter) is designed to work in tandem with the mode switching controller. The controller output, *u* comprises both state-feedback (*−f* (*x̂*, *v̂*)) and an active sensing component (*u_a_*(*t*)). See Methods for details. (b) Triggered Excitation strategy showing the temporal traces of measure of uncertainty (*M_ε_*, trace of the error covariance) with threshold levels *T*_min_ = 4.8 *×* 10*^−^*^3^ and *T*_max_ = 6 *×* 10*^−^*^3^ (top), input *u* (bottom; active sensing component, *u_a_*(*t*): light blue; state-feedback component, *−f* (*x̂*, *v̂*): dark blue). The light purple boxes indicate the triggered time intervals. The triggering started as the trace of the error covariance exceeded *T*_max_ and the triggering continued till *M_ε_* dropped below *T*_min_. (c–e) Simulated position traces (actual states: teal, estimated states: red) using three different exploratory movement strategies for *u_a_*(*t*): Zero Excitation (c), Persistent Excitation, i.e., continuous Gaussian (d), and Triggered Excitation (e) along with the state-feedback. The respective RMS values of the tracking error (*e*_RMS_) are shown in the panels. (f-k) Controller input traces (f–h) and velocity histograms (i–k) for various schemes as in (c–e). Respective RMS values of the inputs (*u*_RMS_) are shown in the panels of (f–h). In (i–k) the fits with a normal distribution (magenta dashed) and three-component Gaussian mixture model (blue solid) are shown along with the respective kurtosis (*κ*) values. (l) Effect on exploratory movement (variance of *u_a_*(*t*)) on tracking error (*e*) and control effort (*u*) in Persistent Excitation. The minimum RMS tracking error and the corresponding control effort are denoted as *e*_PE,min_ and *u*_PE,min_, respectively. (m,n) Effect of threshold pair (*T*_max_, *T*_min_) in Triggered Excitation on tracking error (m) and control effort (n) at optimum variance of *u_a_*(*t*) corresponds to *e*_PE,min_. The solid lines in (m,n) show the respective mean and the shaded regions in (m) correspond to respective SEM (*n* = 300 independent simulations). Note that with the right choice of threshold pair, Triggered Excitation scheme can achieve lower tracking error with substantially lower control effort.

The key feature of the model is that the nonlinear sensory system (i.e. “motion dependent sensor”) embodies the high-pass filtering (i.e., fading or adapting) characteristics found across biological sensory systems.^26, 31–37^ This sensory system model (“motion dependent sensor” in Fig. 5a) assumes nonlinear measurements that decay to zero over time in the face of constant stimuli:

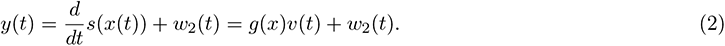

Here, *s*(*x*) is the position dependent sensory stimulus experienced by the organism, *g*(*x*) is the spatial derivative of the sensory stimulus (*ds*(*x*)*/dx*), and *w*_2_(*t*) is the sensory noise. The controller includes a state-feedback-based task level control policy, *−f* (*x̂*, *v̂*), that exploits previously collected sensory information; that information is parsimoniously encoded (i) in estimates of the position and velocity (*x̂*, *v̂*), and (ii) in an ongoing measure of uncertainty, *M_ε_* (based on the covariance of position and velocity estimates). Previous theoretical work has demonstrated that exploratory movements are required for state estimation in control systems that rely on such high pass (i.e., fading) sensors.^24, 25^ Hence, the controller also includes an active sensing control policy, *u_a_*(*t*), that seeks to gain new information through exploratory movements.

To find the optimal balance between exploit and explore components, for a given admissible control policy, *π* and a given weight, *r* for an input, *u ∈ U* (action space), we can define the average steady state cost function:

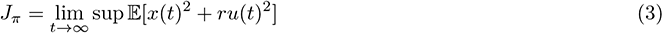

where E is the expectation computed over all the trajectories induced by admissible control policy, *π*. Note that a control policy is admissible if it depends causally on the sensor and actuator data. We chose the cost function, *J_π_* as a weighted combination of steady state tracking error and control effort. Even with complete knowledge of the system states, computation of the optimal solution *J_π_*∗ = inf*_π∈_*_Π_ *J_π_* where Π is the set of all admissible control policies, is only tractable in the case of linear systems or systems with finite state and action spaces.^38^ Since the system is partially observed, existing approaches to optimal control require the solution to an optimal filtering problem and then formulating feedback laws on the filter states.^38^ The filtering problem requires computation of the conditional probability 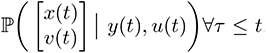. However, due to nonlinearity in the measurement (Eqn. (2)), there is no tractable method to compute this conditional probability, and so heuristic strategies are required. We tested three exploratory movement heuristics for the controller to find an approximate answer to this intractable optimal control problem:

1. **Zero Excitation:** This is a passive strategy (i.e., no exploration) in which the system provides no input excitation for the actuation forces (*u_a_*(*t*) = 0 for all *t*). This is a conventional state-feedback controller.
2. **Persistent Excitation:** This scheme tests a common continuous exploration strategy used in the field of adaptive control.^13^ The controller continually injects a Gaussian input *u_a_*(*t*).
3. **Triggered Excitation:** This mode-switching strategy depends on lower and upper thresholds, *T*_min_ and *T*_max_; the controller only injects Gaussian input when the uncertainty in the state estimator *M_ε_* exceeds *T*_max_, and then continues to inject input until this uncertainty drops below a lower threshold, *T*_min_ (Fig. 5b).

As prior theoretical work shows,^25^ the Zero Excitation strategy (i.e., traditional state-estimate feedback) cannot minimize the state estimation error and hence, not surprisingly, results in poor tracking performance (Fig. 5c), thus illustrating the need for an additional active sensing component in the face of adaptive sensing and perceptual fading. The Persistent Excitation and Triggered Excitation strategies both facilitate substantially better position control than does the Zero Excitation strategy (Fig. 5c,d). Though these two strategies resulted in comparable tracking errors (*e*_RMS_, Fig. 5b,c), the Triggered Excitation was more efficient, requiring substantially lower control effort (*u*_RMS_, Fig. 5g,h). Moreover, unlike the Persistent Excitation strategy, Triggered Excitation generated a distinctive broad-shouldered velocity distribution that featured a sharp peak near zero, with broad shoulders corresponding to bursts of fast movement (Fig. 5j,k). This distribution was strikingly similar to experimental observations across organisms (Fig. 1, Fig. 4, Extended Data Fig. 4, Extended Data Fig. 5, and Extended Data Fig. 6), suggesting that such broad shoulders are a signature (if not definitive proof) of a mode-switching strategy.

We showed that active exploration is essential for better tracking performance as it improves state estimation. But, there is a point of diminishing returns: although higher (more energetic) active excitation can result in excellent state estimation, there is a point beyond which these additional active sensing movements lead to greater tracking errors.

To contrast between Persistent and Triggered Excitaiton we performed a numerical study to obtain the variance of the active sensing signal *u_a_*(*t*) that minimizes RMS tracking error for the Persistent Excitation(PE) strategy, *e*_PE,min_. Note that Persistent Excitation is the limiting case of Triggered Excitation with extremely low threshold values (i.e., insuring that the active sensing mode is always “on”). With that optimum stimulation obtained from Persistent Excitation, we next performed a parameter sweep involving threshold pair (*T*_max_, *T*_min_) in the Triggered Excitation. We discovered that the choice of thresholds in the Triggered Excitation strategy plays an important role—with the right choice of parameters we could achieve better tracking performance (Fig. 5m) at reduced control effort (Fig. 5n). The choice of thresholds also shapes the velocity to best extract sensory information; with low thresholds, the statistics approach that of the Persistent Excitation, whereas high thresholds lead to velocity distributions with higher kurtosis (departure from normality), while requiring less control effort (Extended Data Fig. 7a–c).

How does sensory salience affect performance of the Triggered Excitation (mode-switching) heuristic? To simulate the effects of changes in sensory salience, we parametrically varied the sensory noise variance while keeping constant the switching thresholds *T*_min_ and *T*_max_. As the sensory noise variance was increased (simulating a decrease in salience), the kurtosis value of the velocity distribution decreased, numerically approaching normality in the limit of high sensory noise (Extended Data Fig. 7d–g). This trend of decreased kurtosis in the face of increased noise variance captures the widespread observation that in animals the velocity statistics tend toward a Gaussian distribution as sensory salience is decreased. Moreover, the underlying mechanism, namely increasing frequency of bursts of exploratory movements, matches our experimental observations in *E. virescens*, which performed more frequent transitions to fast movements and spent less time in the slow mode in the lights-off trials than in the lights-on trials (Fig. 2c,d). These analyses clarify that this trend toward a Gaussian distribution with decreased salience is an epiphenomenon of mode-switching: as the frequency of fast movement bursts increases, it overwhelms the task-oriented movements, diminishing the prominence of the central peak.

## Discussion

We examined explore-exploit tradeoffs in the context of goal-direct motor behaviors, such as station keeping, postural balance, and plume tracking, that require active, exploratory movements to enhance sensation. Similar tradeoffs have been extensively studied in categorically different domains, including foraging^33, 39^ and reinforcement learning.^40^ Here we show that the velocity distributions that emerge from the interplay between exploratory movements and goal-directed control are broad-shouldered across taxa, and that this distinctive distribution of movements is robustly modulated by sensory salience. The bouts of ancillary movements that comprise the broad shoulders of these velocity distributions are commonly described as “active sensing”, i.e., the expenditure of energy by organisms for the purpose of sensing^41^*^,a^* . Active sensing research in humans, in relation to touch, was popularized in the 1960’s by J.J. Gibson,^1^ and the original ideas date back at least to the 18th century (for a historical account, see Zweifel *et al.*^41^).

Surprisingly, active sensing is largely avoided in engineering design despite being ubiquitous in animals. The performance of engineered systems may benefit from the generation of movement for improved sensing. An algorithm known as Ergodic Information Harvesting (EIH)^3^ could be used to control movements for sensing in artificial systems. This algorithm balances the energetic costs of generating movements against the expected reduction in sensory entropy. The EIH has been tested in relation to several animal model systems, and produces plausible animal trajectories.^3^

Interestingly, the EIH algorithm produces the opposite trend in kurtosis of velocity distributions in relation to sensory salience (Extended Data Fig. 2g–l, Extended Data Table. 3) that we observed in our experiments, reanalysis of prior data, and in our model: as sensory salience decreases, there is an increase in active sensing movements but a decrease in kurtosis (Extended Data Table. 2). That EIH leads to decrease in kurtosis occurs in part because EIH generates continuous sensing movements, and does not incorporate mode switching. A refined EIH model, that generates the temporally distinct periods of sensing movements that characterize mode switching, would better reflect our findings in animals, and is a promising strategy for improving the performance of robotic control systems. How mode switching is manifest across the diverse biological systems we examined is a compelling open question. Many of these control systems have evolved via convergent evolution in which adaptive strategies emerge independently across lineages. One result of convergent adaptation is that species often rely on idiosyncratic features, such as feathers or skin flaps, to achieve the same adaptive strategy, such as flapping flight. We infer that the mechanisms for mode switching is present in control systems that range from sub-cellular systems^47^ to neural systems in vertebrates.

The mechanisms for mode switching in vertebrate nervous systems may emerge at different levels within sensorimotor control pathways. For example, neurophysiological recordings show that sensory salience can be encoded in brain circuits via synchronization and desynchronization of spiking activity.^48^ Such population coding of salience,^49–51^ when coupled with a threshold, could trigger discrete bursts of motor activity for sensing.^8^ Motor circuits for the production of discrete bursts of movement occur in spinal circuits.^52^

These discrete bursts of movements could arise from reflex-like, threshold-based activity in animals, akin to how Mauthner cells trigger a cascade of motor activity when sensory inputs exceed a threshold.^53^ A key difference between reflex-like, threshold-based behaviors and the mode-switching we describe in this paper would be that the signal in question would arise from an internal representation of sensory uncertainty, rather than from the overall level of sensory excitation. Such a reflex-like action could produce stereotyped forms of interactions with the external environment in relation to sensing.^8^

A common engineering approach to sensing and control is to add sensors and improve sensor performance, particularly at low frequencies, effectively side-stepping the need for active sensing movements altogether. Such improved sensing enhances observability without relying on movement. In stark contrast, organismal sensor systems are almost invariably adapting (high-pass), necessitating active sensing. Irrespective of whether organisms have achieved an optimal solution to the control problem (or instead are limited by evolutionary constraints on sensor performance), the widespread convergent evolution of a common active sensing strategy nevertheless suggests an alternate engineering design paradigm. The confluence of adapting sensors^54^ and the uncertainty-triggered mode-switching heuristic presented in this paper provides a new roadmap for movement control of robotic systems.

## Methods

### Tracking of glass knifefish

#### Subjects

We obtained adult, weakly electric, glass knifefish *Eigenmannia virescens* (10 – 15 cm in length) from commercial vendors, and housed the fish according to the published guidelines.^55^ Water temperature in the experimental tank was kept between 24 – 27 *^◦^*C, and conductivity ranged from 10 – 150 µS*/*cm. Fishes were transferred from the holding tank to the experimental tank 12 – 24 hours prior to the experiments, to allow for acclimation. All experimental procedures were approved by the Johns Hopkins Animal Care and Use Committee, and followed guidelines established by the National Research Council and the Society for Neuroscience.

#### Experimental apparatus

The experimental apparatus was similar to that used in previous studies.^2, 6, 8, 23, 56^ The refuge was machined from a 152 mm long segment of 46 *×* 50 mm rectangular PVC tubing, with the bottom surface removed to allow the camera to record the ventral view of the fish. On both sides of the refuge, a series of six rectangular windows (6 mm wide *×* 31 mm high, spaced 19 mm apart) were machined, through which to provide visual and electrosensory cues.

A computer sent designed digitized input stimuli (25 Hz) from LabVIEW (National Instruments, Austin, TX, USA) to a FPGA based controller for a stepper motor (STS-0620-R, HW W Technologies, Inc., Valencia, CA, USA). The stepper motor drove a linear actuator, leading to the one degree of freedom refuge movement in real time. A video camera (pco.1200, PCO AG, Kelheim, Germany) captured fish movements through mirror reflection at 100 Hz. The captured frames (width × height: 1280 pixels *×* 276 pixels) were saved as 16 bits .tif files via camera application software (pco.camware, PCO AG, Kelheim, Germany).

#### Experimental procedure

The experiments were conducted in two illuminance levels—around 0.3 lx (lights off) and 80 lx (lights on). Each trial lasted for 60 s. During the initial 10 s of each trial, the refuge was actuated to follow a 0.45 Hz sinusoidal trajectory, the amplitude of which was gradually increased to 3 cm, and then decreased to 0 at the end of the 10 s interval, in a similar fashion as described in Biswas *et al.*^2^ After the initiation phase, the refuge remained stationary for 40 s, finally followed by a termination phase for 10 s, during which the refuge was actuated in a similar fashion as during the initiation phase.

#### Tracking Algorithm

To observe fine details of the fish movement, we used a high frame rate in our video recordings. High tracking accuracy was essential as the position and velocity data were likely to be contaminated by measurement noise. To ensure high tracking accuracy, the refuge and fish position were analyzed by custom video tracking software^57^ developed by Baĺazs P. Vágvölgyi from the Laboratory for Computational Sensing and Robotics (LCSR), at Johns Hopkins University.

The tracking algorithm worked in two phases. The first phase was template matching, which roughly located the targets (fish or refuge). In the first frame of a given video, a rectangular region was manually selected around the target to create a template. On subsequent frames, a neighborhood region around the template (*±* 20 pixels) from the previous frame was selected for the computation of a normalized 2-D cross-correlation matrix. If the target changed its orientation in the new frames, before computing the normalized 2-D cross-correlation, the new image frame was first rotated to match the orientation of the template from previous frame. If needed, the areas of the image were sampled (then scaled and interpolated if necessary) with subpixel accuracy.

After creating the template, the second phase applied the Levenberg-Marquardt algorithm to find the global maximum of the normalized cross-correlation function. This step produced a match between the template and target at each frame, with subpixel accuracy. We performed extensive preliminary testing and analysis to confirm that the remaining measurement errors had smaller variance than the stochastic movements of the fish.

#### Data Processing

The tracking algorithm stored the fish position in both horizontal and vertical directions, (originally in pixels, along with the respective pixel to meter conversion factor), and the angle of orientation (in degree) in .csv files. We used only the data while the refuge was stationary (40 s, 4000 data points in total) for each trial. To further reduce the measurement noise, the position data was filtered through a Butterworth zero-phase distortion filter (filtfilt command in MATLAB) with a 15 Hz cutoff frequency. Fish velocity in the horizontal direction was computed as forward differences of the horizontal position time series.

#### Identification and Characterization of Behavioral Modes

For the identification of the behavioral modes, we used three different clustering approaches—(1) Gaussian mixture model (GMM) with inflection point based clustering, (2) hidden Markov model (HMM) based clustering, and (3) GMM with maximum a posteriori probability (MAP) based clustering.

For GMM with inflection point based clustering, the velocity (*v*) data from each individual fish at a specific lighting condition were clustered into three components, slow, fast positive, and fast negative, using two velocity thresholds, *v_L_* and *v_H_* (*v_L_ < v_H_*), resulting in two behavioral modes—slow and fast (fast positive and negative were combined). The velocity threshold values were computed by finding the inflection points of the GMM fits to the velocity data, *f*_GMM_, specific to a lighting condition. To numerically identify the inflection points of *f*_GMM_, we numerically computed the spatial second-order derivative of *f*_GMM_’’ (*f*), and located the first and the last indices of the array 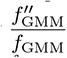 such that the condition 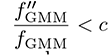 was satisfied for a given *c*. This method separated the central peak of the *f*_GMM_, velocity distribution around zero velocity, from the broad shoulders. We chose *c* = 0.005 for all the individual fish irrespective of the lighting conditions, except for fish 1 lights-off trials (*c* = 0.02), (the different c value for fish 1 lights-off trials was chosen so that the relative area under the central peak of the distribution was less than 0.6 (similar to other fish during lights-off trials). For further analysis of these behavioral modes, we assumed a continuous time Markov chain model. For infinitesimal *dt*, the transition probabilities, *Q_ij_* are given as follows:

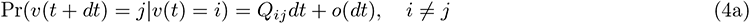

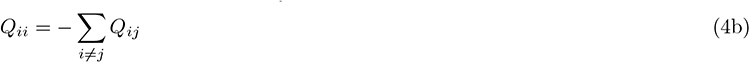

and the probability matrix **P** with *p_ij_* = Pr(*v*(*t*) = *j|v*(0) = *i*) and transition rate matrix **Q** with entries *Q_ij_* satisfy the first order differential equation

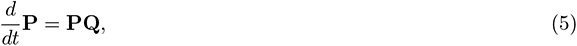

whose solution is given by,

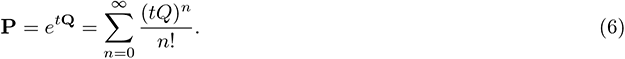

For every trial from each individual fish, we computed the probability matrix **P** with entries *p_ij_, i* = 1, 2*, j* = 1, 2 where state 1 and 2 correspond to slow (exploit) and fast (explore) modes, respectively. We used the approximation to the matrix exponential in Eqn. (6), 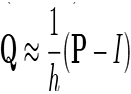 for the computation of the transition rates between slow and fast modes in each trial from the respective probability matrix, **P**. All analysis was performed using custom code written in MATLAB.

For HMM clustering, we combined all the positional trial data (*x_t_*) from all the five fish at a specific lighting condition along with their negatives (*−x_t_*). We included the negative data to get rid of any directional bias. Using the NHMSAR package in R we fitted a homogeneous Markov switching first-order autoregressive model with three hidden states and Gaussian innovation to the position data as follows:

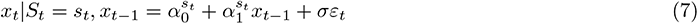

where, *x_t_* is the observed position, *S_t_* is the hidden univariate process defined on *{*1, 2, 3*}*, and *ε* is Gaussian white noise. The HMM fitting resulted in three clusters similar to slow, fast positive, and fast negative as obtained with GMM with inflection point based clustering method. Finally by combining fast positive and negative, we ended up with two behavioral modes—fast and slow for further computation of switching frequency and residence time.

In GMM with MAP based clustering, GMM models with three components were fitted to the velocity data from each individual fish at a specific lighting condition. We assigned the cluster index for each data point based on the maximum a posteriori probability using the Bayes’ rule. We found a careful inspection is always necessary for this clustering approach as the central “exploit” peak of the velocity distributions was, in some cases, captured by more than one Gaussian mixture component. For representative data from fish 1, we employed two methods. In Method 1, we treated all the three clusters separately for the computation of residence time. In Method 2, we combined the two clusters associated with Gaussian mixture components with lower variances into single slow mode. All the analysis was performed using custom code written in R and MATLAB.

### Simulation

Sensory adaptation is a robustly observed phenomenon among organisms ranging from unicellular amoebae^36, 37^ to humans^26^ where the sensory systems stop responding to constant stimuli. Here, we modeled this adaptive/ high-pass nature of the sensory receptors as a “motion dependent sensor” for which we assumed a nonlinear measurement model^24, 25^ with sensory noise *w*_2_(*t*):

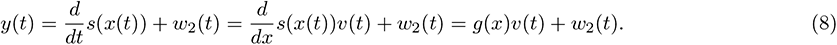

Here, *s*(*x*) is the position dependent sensory scene experienced by the organism. For the present study, we assumed a quadratic sensory scene *s*(*x*) = 1/2 *αx*^2^ +*βx* with nonzero constant sensory scene parameters *α* and *β*. This assumption on sensory scene yields *g*(*x*) = *αx* + *β*, a linear function of position, *x*.

Due to the presence of the nonlinearity in the measurement, we used an extended Kalman filter (EKF) for state estimation, a common heuristic. For the state-feedback component we applied *f* (*x̂*, *v̂*) = *k*_1_*x̂* + *k*_2_*v̂*. In the Triggered Excitation scheme, for the uncertainty measure (*M_ε_*) we used the trace of the state estimation error covariance matrix, Tr(*P* (*t*)). When the uncertainty measure Tr(*P* (*t*)) rose above a maximum threshold, *T*_max_, the controller generated active sensing component, *u_a_*(*t*) as a Gaussian input with fixed power spectral density and it continued to inject the input until Tr(*P* (*t*)) dropped below a lower threshold, *T*_min_. At this point the controller switched back to traditional state-feedback form. For the Persistent Excitation scheme, the controller continued to inject a Gaussian input *u_a_*(*t*) for all time.

To obtain the critical excitation level of the active sensing component *u_a,_*_crit_(*t*) for optimum tracking performance in Persistent Excitation, we chose 30 logarithmically spaced variance values of *u_a_*(*t*) from 1 to 100. From the mean of 100 independent simulations for each variance value, we obtained *u_a,_*_crit_(*t*) *≈* 9.33 which achieved the minimum RMS tracking error of *e*_PE,min_ *≈* 0.071 and RMS control effort *u*_PE_ *≈* 10. Using this critical value for the excitation *u_a,_*_crit_(*t*), we studied the effect of thresholds in Triggered Excitation by varying *T*_max_ and *T*_max_*/T*_min_ linearly from 4 *×* 10*^−^*^3^ to 10 *×* 10*^−^*^3^, and 0.5 to 1, respectively, and performed 300 independent simulations for each pair of values.

The system parameters were chosen from prior studies^6, 24^ as follows: *b* = 1.7, *m* = 1, *α* = 3, *β* = 5, *k*_1_ = *mω*_n_^2^, *k*_2_ = (2*mζω_n_* + *b*), *ζ* = 0.56, *ω_n_* = 1.05 *×* 2*π*. The process noise, *w*_1_(*t*) and sensor noise, *w*_2_(*t*) were chosen as fixed Gaussian noise inputs with variances 0.03 and 10, respectively.

### Statistics

All the statistical analysis was performed with sign test and Mann-Whitney-Wilcoxon (MWW) test using custom codes written in R (R core team, Indianapolis, Indiana, USA) and MATLAB (Mathworks, Natick, Massachusetts, USA). For all tests, the significance level was set to 0.05. The experimental and simulation data are provided as either mean plus or minus the standard deviation (*µ±*SD) or mean plus or minus the standard error of the mean (*µ±*SEM).

## Data and software availability

An archived version of the datasets and the analysis code supporting this article will be made available through the Johns Hopkins University Data Archive and on Code Ocean. All the figure data will be made available through figshare.

## Acknowledgement

We thank Tim Kiemel (UMD) for providing human balance data and Baĺazs P. Vágvölgyi (JHU) for developing the tracking software used in this work. We thank Cynthia F. Moss (JHU), Varun P. Sharma (GT) and Simon Sponberg (GT) for suggesting relevant articles for the reanalysis of the animal locomotion data, and Sarah L. Poynton (JHMI) for critical feedback on the manuscript. This work was supported by Office of Naval Research under grant no. N00014-21-1-2431 (N.J.C).

## Author contribution

Conceptualization, all authors; Methodology, all authors; Software, D.B., A.L., K.H., Y.Y. and J.G.; Formal Analysis, D.B., A.L., K.H., Y.Y. and J.G.; Investigation, Y.Y.; Data Curation, D.B. and Y.Y.; Writing – Original Draft, all authors; Writing – Review and Editing, all authors; Visualization, D.B. with other authors; Supervision, N.J.C.

## Supplementary Materials and Methods

All experimental data for reanalysis were acquired from prior studies either directly through the authors^17^ or published dataset.^14–16, 18–20, 22^ All computations for reanalysis were performed using custom R and MATLAB scripts.

### Refuge tracking of glass knifefish in presence of jamming electrodes

We examined experimental data for refuge tracking of *Eigenmannia virescens* under varying electrosensory conditions from a prior study.^3^ In that study, a rectangular refuge was actuated to follow a sinusoidal trajectory of frequency 0.1 Hz and amplitude of 17 mm (in contrast to the stationary refuge in our study). The fish position was tracked at 60 Hz. The experiments were conducted in the dark, in the presence of a closed-loop jamming system, where the frequency of the jamming signals was set to 5 Hz below the electric organ discharge (EOD) frequency of the fish.

To extract the active exploratory movements from the refuge tracking response, we computed the Fourier transform of the temporal response of the fish for a given trial, deleted the frequency peak corresponding to the refuge motion (0.1 Hz) from the frequency response, and then reconstructed the time domain signal using the inverse Fourier transform. To estimate the velocity of the active exploratory component, we scaled the forward difference of position data.

### Postural sway in humans

Experimental data of human postural sway in upright stance were acquired from a prior study^17^ through the authors. In the original experiment, nine healthy adults (subjects) maintained a modified tandem Romberg stance for 4 min. Trials were performed under one of four multi-sensory conditions, with different combinations of vision and tactile sensing: (i) neither vision nor touch, (ii) only vision, (iii) only touch, and (iv) both vision and touch. Data for the mediolateral center of mass (ML CoM) were collected from a sensor attached to each subject’s lower back (at the level of the fourth or fifth lumbar vertebra).

In the present study, we analyzed conditions ii, iii, and iv in which the subjects used vision and/or tactile cues. From the original study, we excluded the data from two subjects as the velocity data from them could not be grouped into a unimodal distribution, unlike the data from other subjects. Thus we ended up using *N* = 7. The original data were collected at a frame rate of 50 Hz. We subsampled the data at 5 Hz to reduce the noise in the velocity data, which was computed from forward differences. For the identification of the “explore” and “exploit” behavioral modes, we used GMM model with inflection point based clustering and computed the switching frequency among modes.

### Saccadic eye movements in humans

Data for eye movements in humans during fixated gaze at a stationary object were acquired from a prior study involving five adult humans (subjects) with normal or corrected-to-normal vision.^21^ The subjects were instructed to look at circular stimuli of different levels of Gaussian blur, and matched summed intensities on the screen, at a distance of 175 cm. The eye movements of the subjects were recorded using an Eyelink 1000 eye tracker. The original data were collected at 1000 Hz, then smoothed by the previous authors using a third-order Savitzky-Golay filter of frame length 21 ms. The prior authors also removed blink periods, as well as the data 200 ms before and 300 ms after each blink.

In the velocity data we detected and removed outliers (velocity value greater than 1.5*×* degree/ms) that were likely caused by measurement errors; these outliers comprised only 0.05% of the original data.

### Bilateral eye movements in house mice

Data for eye movements in freely moving house mice, *Mus musculus*, during capture of prey (crickets) in a closed arena, were acquired from a prior study.^15^ Eye and head movements were classified by the original authors into two different behavioral phases: (i) approach and (ii) non-approach. During the approach phase, the head of the mouse was oriented towards the cricket (the azimuthal angle was within *±*45 degree), and the mouse was locomoting faster than 1cm*/*s. During this approach phase, the authors reported that the left and right eye movements were more synchronized with each other than in the non-approach phase.

The approach phase behaviors were short (*≈* 5.8% of the whole dataset), whereas non-approach segments were long and more frequent. Both during the non-approach and approach phases, we observed typical broad-shouldered velocity distributions with kurtosis (*κ*) greater than 3, although mice exhibited less exploratory behavior during approach phases (*κ* = 10.37 and 8.56 for right eye and left eye, respectively). In our present study, we focused on the data during the non-approach phases. The original dataset had 105 trajectories collected from seven mice at a frame rate 30 Hz. The data contained some discontinuities due to tracking loss. To fill those missing data, we used a 30-sample moving median filter. The eye velocity data was computed using forward differences.

### Pinnae movements in big brown bats

Echolocation data were acquired from a prior study^16^ of pinnae movement in big brown bats, *Eptesicus fuscus*, in relation to prey trajectories. The prey targets were mealworms tethered to a wire which were moved from an initial distance of 2.5 m toward the bats. Bats were trained to rest on a platform and track the target using echolocation. For our current analysis, we considered two of the four conditions originally presented:^16^ (i) single-target simple-trajectory (Fig. 5d, Supplementary Fig. 5b–f, left), and (ii) single-target complex-trajectory (Supplementary Fig. 5b–f, right). In the former scenario, there was only forward movement of the target toward the bats; in the latter case the target was receded twice before finally arriving at the bat platform. The dataset had 10 trajectories for each condition collected from three bats at frame rate 500 Hz. The echolocation calls of the bats were classified into “approach” and “buzz” phases.^58^ The approach phase was characterized by the relatively constant and higher pulse duration and pulse interval, whereas the buzz phase was characterized by a sharp drop in both. A distance threshold of 0.5 m between the mealworms and the bats roughly correlated with the onset of the buzz phase (Fig. 1B,C in^16^). We focused on examining the data during the approach phase only. From the *z*-positions of the tip of the pinnae, we computed the relative pinnae movement, *Z_L−R_* as the difference between *z*-positions of left and right pinnae tips. From this we then computed the rate of change of this relative movement using forward differences in *Z_L−R_*.

### Analysis of odor tracking in Eastern moles

Blind Eastern moles *Scalopus aquaticus* are known for their excellent odor tracking capability. We took the tracking data of moles locating a stationary odor source (small sections of earthworms), in a semicircular arena, from a published dataset,^3^ which was digitized from the original study.^14^ Among the three original experimental conditions, we only considered the case of normal airflow where the moles had their two nostrils open. The dataset comprises six trajectories from two moles. To align the data with respect to the destination, we subtracted the last datapoint from the angular position data. Although the frame rate for data collection was not stated in the original study, it was reported that the moles usually found the odor sources in 5 – 10 s, and the longest data sequence had 125 frames. Thus, we set the frame rate at 25 frames per second when computing the angular velocity using forward differences; note that an error in frame rate would not affect our results as it only effects the scale of the velocities.

### Odor plume tracking in American cockroach

Plume tracking data of adult male American cockroaches, *Periplaneta americana*, were acquired from a published dataset^3^ which derived from a prior study.^18^ The head position was tracked in males during their tracking task of locating the source of female sex pheromone (Periplanone B) in a laminar flow tunnel (with wind speed 25 cm*/*s) under red and infrared light. To understand the role of the olfactory sensors on the antennae, the authors conducted experiments in which either one antenna (unilateral) or both antennae (bilateral) were cut to specific lengths. The original data were collected at a frame rate of 30 Hz, which was later subsampled by the original authors at 15 Hz. In the present study, we only considered lateral movement of the head for cockroaches in which there had been bilateral shortening of the antennae. For the identification of the “explore” and “exploit” behavioral modes, we used GMM model with MAP based clustering and computed the respective residence times. The statistical test between antennae length “4 cm” and “2 cm” resulted p-values greater than significance level 0.05, although the trend was certainly in the anticipated direction—with decrease in sensory salience RMS movement increases and residence time at slow mode decreases.

### Analysis of tactile sensing in Hawkmoths

Hawkmoths (Sphingidae) use their proboscis to actively explore the flower surface in search of the nectary. We acquired the tactile sensing data for the Carolina sphinx hawkmoth, *Manduca sexta* from a prior study^20^ through the published dataset. In the original experiment, the moths were presented with 3-D printed flowers (modeled after the corolla), with four different cross-sectional curvature profiles. We analyzed data for three profiles, “funnel”, “near-funnel”, and “near-flat”, for the moth’s first and seventh visit to the flowers. In the experiment, the tip and base of the unfurled proboscis, and the antennae were tracked at a frame rate of 100 Hz. The prior study reported two parameters: (i) normalized relative radial distance (RRD), which was computed as the distance from the proboscis tip to the center of the flower normalized to the radius of the flower; and (ii) the relative radial orientation angle (RRA) angle, which was computed as the angle between the proboscis tip trajectory and radial axis which was further wrapped to restrict the range to 0 – 90 degree.

On successful insertion of the proboscis, the sharp decrease in the RRD indicated that the moth had started feeding from the nectary. Based on this observation, we divided each trial into two phases: pre-feeding and feeding, where the transition frame was manually selected by inspection of the data. We only used the tracking data pertinent to the pre-feeding phase. We discarded the trials where these two phases were not distinct. We computed the rate of change of RRA (*ω_RRA_*) from forward differences.

### Analysis of visual tracking of flower by Hawkmoths

Hawkmoths rely heavily on visual inputs to track a swaying flower, and hover during tracking, with lighting conditions affecting tracking behavior.^19^ We analyzed tracking data published in an earlier study.^19^ In the original experiment, three species of hawkmoths (nocturnal *Deilephila elpenor*, diurnal *Macroglossum stellatarum*, and crepuscular *Man-duca sexta*) were presented with a 3-D printed flower, actuated by a stepper motor to follow a sum-of-sine stimulus. The experiments were performed at three different illuminance levels: 300, 15, 0.3 lx for *D. elpenor* ; 3000, 300, 15 lx for *M. stellatarum*; 300, 15, 0.3 lx for *M. sexta*. The datasets were collected at 100 Hz.

For our present analysis, we considered the data for 15 and 300 lx being the common lux levels tested across all three species of hawkmoths. To extract the active exploratory movements from the tracking response, we first computed the Fourier transform of the original temporal response, then removed the frequency peaks corresponding to the flower motion from the frequency response, and finally reconstructed the time domain signal using an inverse Fourier transform. The velocity was estimated from forward differences of position data.

### Analysis of galvanotaxis in amoebae

Amoebae have served as a classic model for studying the movement of eukaryotic cells in response to various chemical, mechanical, and electrical cues. Galvanotaxis (movement in response to electric field) data of movement of two related unicellular organism *Amoeba proteus* and *Metamoeba leningradensis* were acquired from a prior study^22^ through the published dataset. The experiments were performed in batches (experimental replicates). Each time a small group of amoebae was placed in a custom-made chamber with simplified Chalkley medium: *A. proteus* (experimental replicates: 8, number of cells per replicate: 4–8), *M. leningradensis* (experimental replicates: 4, number of cells per replicate: 4–15). Crowding of cells was avoided for ease in tracking individual cell as well as to eliminate any coordinated interaction among cells. Once a laminar flux was established in the experimental chamber, an electric field was applied with a gradient of 334–342 mV*/*mm to study the galvanotaxis. The original data comprising trajectories of 50 cells of each species, were collected at 0.1 Hz.

In the present study we focused on the movement in the transverse direction of the applied electric field. The velocity was estimated from forward differences of position data.

---

**Extended Data Fig. 1.**
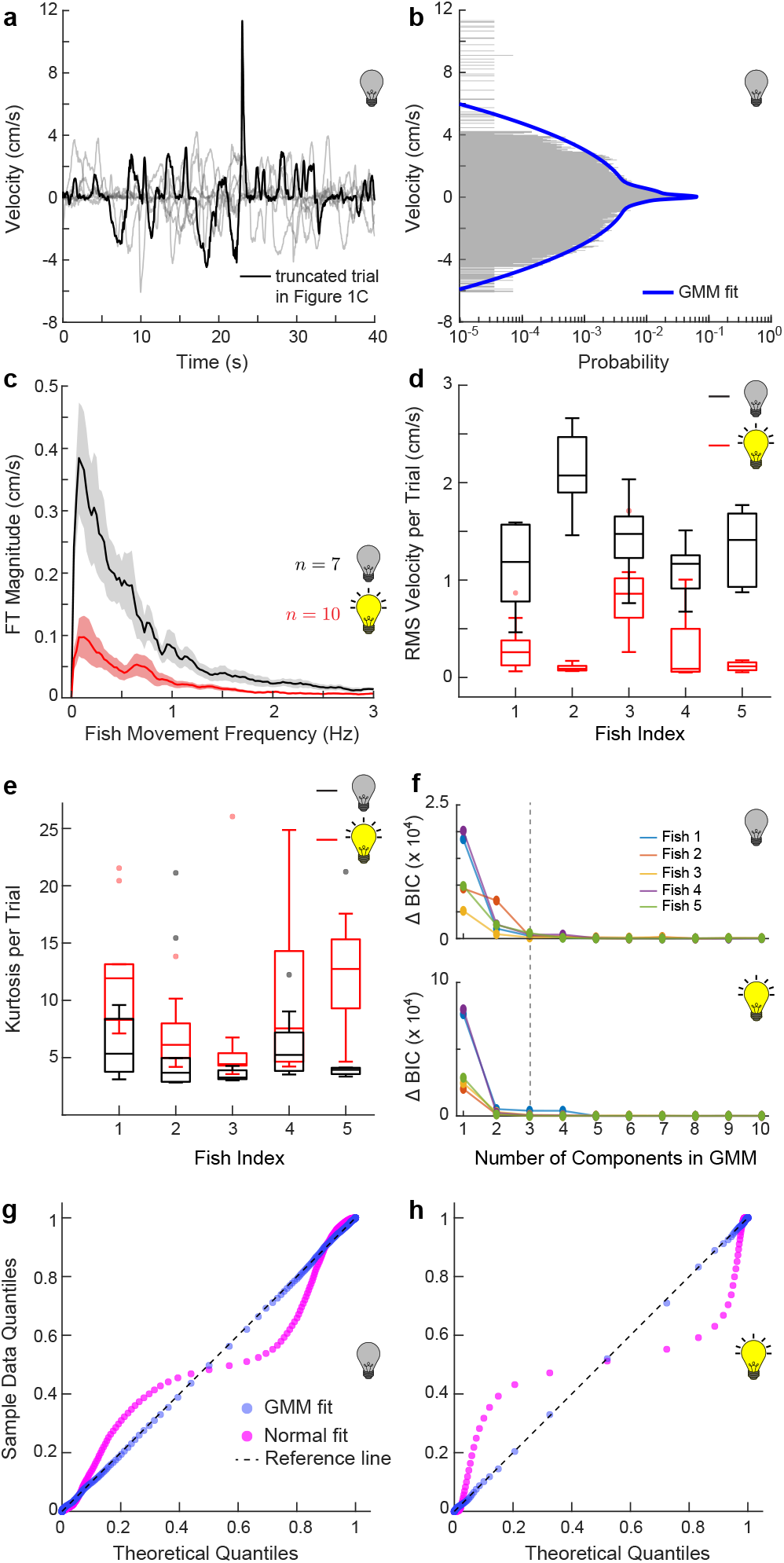
Comparison of fish movement in lights-on trials versus lights-off trials. (a) Velocity trace of the truncated trial data (black) with traces from other trials (gray) for the same fish from Fig. 1c. (b) Histogram of all trials (*n* = 7) using same length scale, and the three-component Gaussian mixture model (GMM) fit. (c) Magnitude of discrete Fourier transform of velocity traces with total number of trials (*n*) indicated next to the plots. The solid line and the shaded region denote mean and the standard error of mean, respectively. (d) Box and whisker plot showing RMS values of individual trials for all fish (*N* = 5) with colors the same as in (c). Mean RMS velocity across trials for all fish in lights-off trials was greater than in lights-on trials (one-tailed p-value *<* 0.005). (e) Box and whisker plot showing kurtosis values of individual trials for all fish (*N* = 5) with colors the same as in (c). For all fish, mean kurtosis values across trials in lights-on trials was greater than in lights-off trials (one-tailed p-value *<* 0.05). (f) Cumulative difference in Bayesian information criterion (ΔBIC) values for varying number of components in GMM (top: lights off; bottom: lights on). The gray dashed line corresponds to three-component GMM. All p-values were calculated using the Mann-Whitney-Wilcoxon test. (g, h) Q-Q plots from a single representative fish (fish 1) comparing the velocity data from lights-off (g) and lights-on trials (h) with theoretical quantiles from a normal fit (magenta) and GMM fit (blue), respectively. Lesser deviation from the reference line (black dashed) for GMM fits with three components than for the normal, suggested better fitting of the former. The Q-Q plots also showed that the lights-off trial data were closer to normal distribution fits than were those of lights-on trial data.

**Extended Data Fig. 2.**
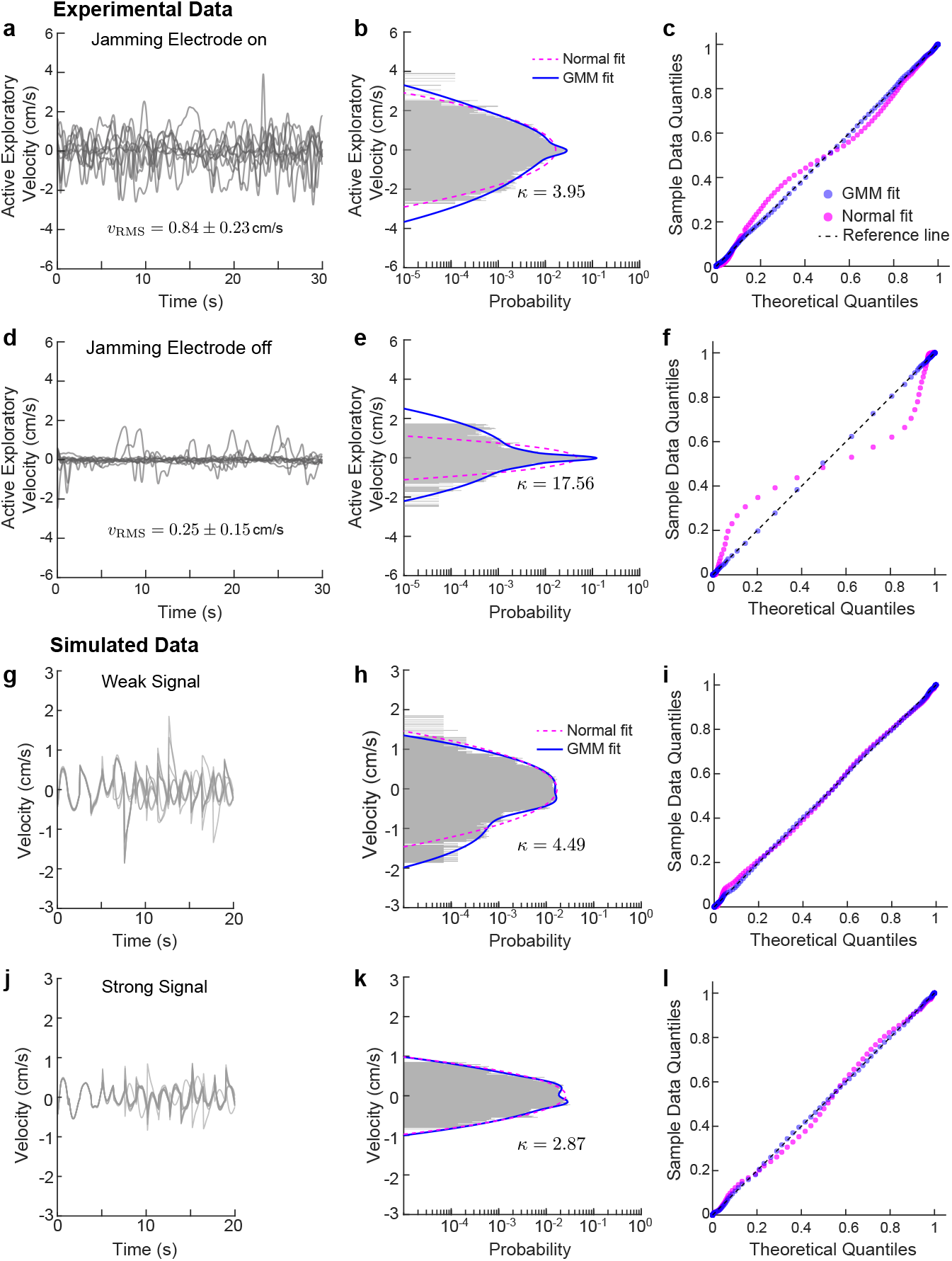
Reanalysis of the experimental and simulated trajectories from Chen *et al.*^3^. (a) Experimental velocity traces (*n* = 10) with “jamming” of the electrosensory system in *Eigenmannia virescens*, which decreased the salience and reliability of electrosensory navigation. (b) Corresponding histogram, with the kurtosis value, *κ*. The magenta dashed and the blue solid curves correspond to a normal and GMM fit with three components, respectively. (c) Q-Q plots comparing the sample velocity data for all trials (*n* = 10) from (a) with theoretical quantiles from the same normal (magenta) and GMM fit (blue) from (b). Clearly, the GMM fit was better than the normal. See Extended Data Table 2 for statistical details. (d–f) Experimental velocity traces (*n* = 10) with jamming electrode off (d), corresponding histogram (e) with the kurtosis value, *κ* and the Q-Q plots (f). (g–i) Simulated velocity traces (*n* = 9) using Ergodic Harvesting Information (EIH) algorithm for weak signal (g; SNR *≤* 30 dB equivalent to jamming amplitude *≥* 10 mA) (g), corresponding histogram (h) with the kurtosis value, *κ* and the Q-Q plots (i). (j–l) Simulated velocity traces (*n* = 9) using Ergodic Harvesting Information (EIH) algorithm for strong signal (d; SNR *≥* 50 dB equivalent to absence of jamming) (j), corresponding histogram (k) with the kurtosis value, *κ* and the Q-Q plots (l). Colors and styles are same as in (a–c). Mean *±* SD of the respective RMS values of experimental velocity traces (*v*_RMS_) are shown next to plots of (a,d). The fitting performances of GMM and the normal distribution for simulated trajectories were comparable. See Extended Data Table 3 for statistical details.

**Extended Data Fig. 3.**
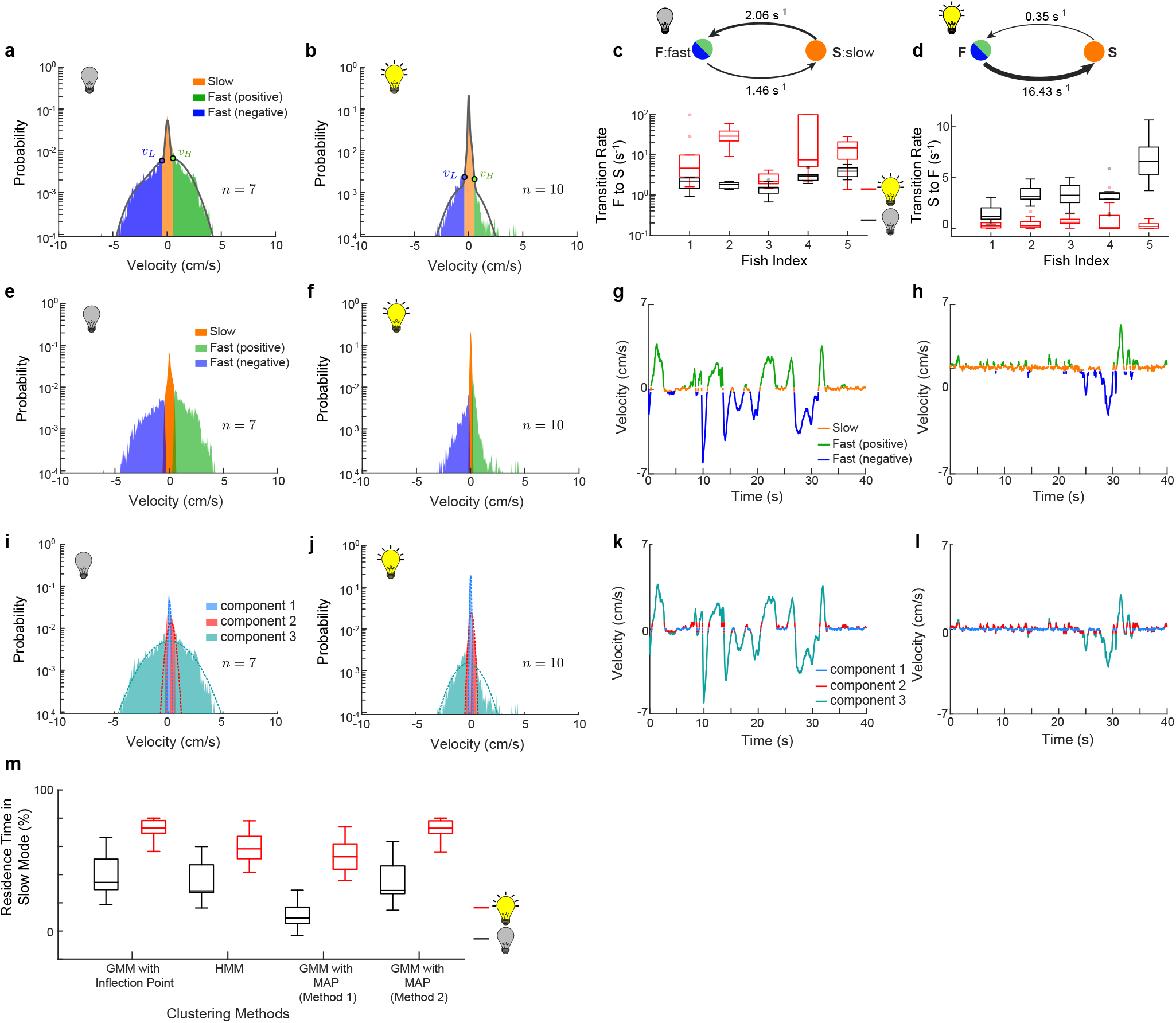
Clustering of velocity data into different behavioral modes. (a,b) Representative velocity histograms of lights-off (a) and lights-on (b) trials from the same fish with three clusters: slow (orange), fast positive (green) and fast negative (blue). The clustering was based on identifying velocity thresholds *v_L_* and *v_H_* on the Gaussian mixture model (GMM) fit (grey line) as indicated by blue and green markers, respectively (see Methods for details). (c,d) Top: two-state (F: fast velocity both positive and negative combined and S: slow velocity) Markov process showing mean transition rates for a representative individual in lights-off (a) and lights-on trials (b). Bottom: transition rates corresponding to state transitions: F *→* S in (e) and S *→* F in (f), respectively for lights-off (black) and lights-on (red) trials. The transition rate for F *→* S was higher in lights-on trials (one-tailed p-value *≤* 0.02) whereas the rate for S *→* F was higher in lights-off trials (one-tailed p-value *<* 0.003). (e–h) Velocity histograms (e,f) and traces (g,h) from lights-off (e,g) and lights-on (f,h) trials from the same fish from (a,b) showing three clusters using Hidden Markov model (HMM) based clustering. The colors are same as in (a,b). (i–l) Velocity histograms (i,j) and traces (k,l) from lights-off (i,k) and lights-on (j,l) trials from the same fish from (a,b) showing three clusters using maximum a posteriori (MAP) clustering based on three-component GMM fits. The probability density functions (pdf) of respective components are shown as dashed lines in (i,j). (m) Box and whisker plots showing residence time in slow mode, computed as the percent of the trial duration (40 s), for lights-off (black) and lights-on (red) trials computed using different clustering algorithm. For details see Methods. For all the clustering algorithms, the computed residence time was significantly higher during lights-on trials than lights-off trials (one sided p-value *<* 0.005). All p-values were calculated using the Mann-Whitney-Wilcoxon test.

**Extended Data Fig. 4.**
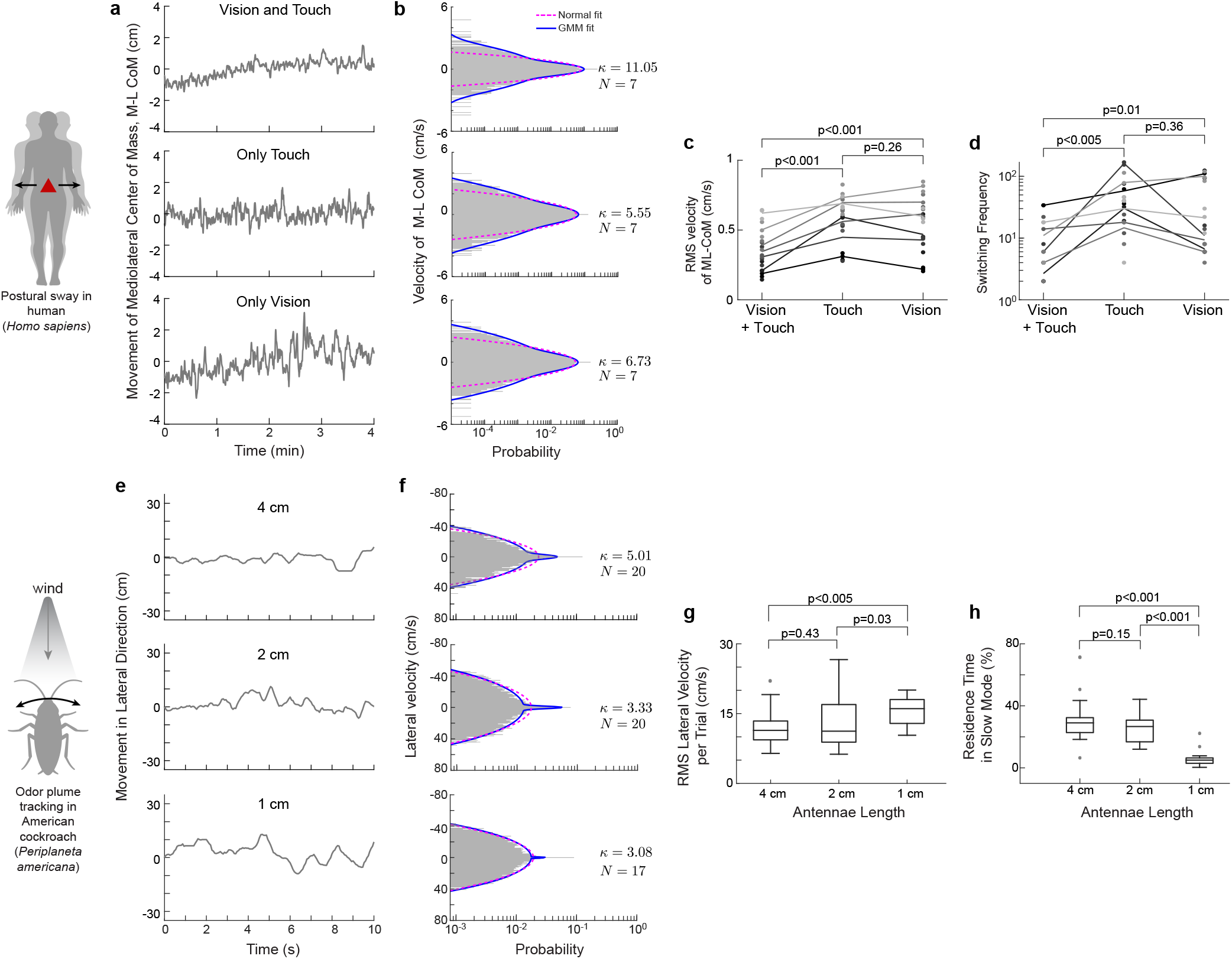
Reanalysis of the postural sway in humans, *Homo sapiens*,^17^ and odor plume tracking response of American cockroach, *Periplaneta americana* ^18^ show evidence in support of sensory salience dependent mode-switching strategy. (a,b) Representative temporal traces of mediolateral movement of center of mass, ML CoM (a) and the histograms of ML CoM velocities (b) for different experimental conditions—both vision and touch (top), only touch (middle) and only vision (bottom). The magenta dashed and the blue solid curves in (b) correspond to a normal and GMM fit with three components, respectively. The dataset analyzed here, comprised of 7 subjects, with three replicate trials per experimental condition, was collected at 50 Hz. (c,d) Comparison of the RMS velocities (c) and switching frequency (d) for different experimental conditions. Different shades of gray denotes different human subjects. The p-values were computed using the sign test. (e,f) Representative temporal traces of the lateral head movement (e) and the histograms of the lateral velocities (f) for different antennae length as indicated. The colors of fitted curves are same as in (a,b). The dataset analyzed here, was collected at 30 Hz but later was subsampled at 15 Hz by the original study authors. The kurtosis (*κ*) values and the total number of trajectories (*N*) analyzed are indicated next to the respective panels in (f). (g,h) Comparison of the RMS lateral velocities (g) and residence time at slow mode (h) for different experimental conditions. The one-tailed p-values were computed using the Mann-Whitney-Wilcoxon test.

**Extended Data Fig. 5.**
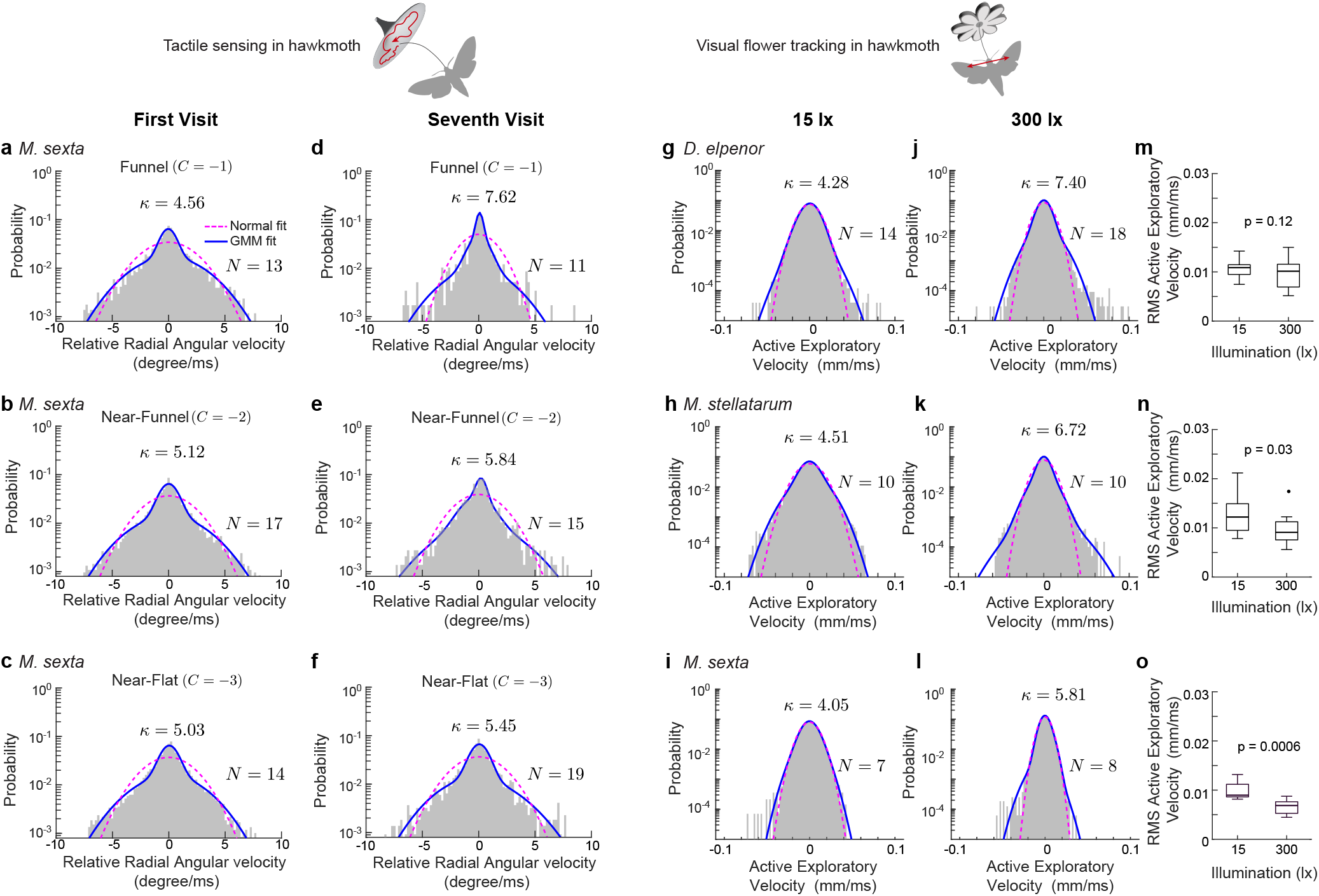
Reanalysis of the tactile response in crepuscular hawkmoth, *Manduca sexta*,^20^ and active exploratory movement of three different species of hawkmoths during flower tracking^19^ shows similar broad-shouldered velocity distributions. (a–f) Histograms of relative radial angular velocity for different shape of the flower as indicated. “*C*” is the curvature parameter for the description of the lateral traces of the corollas for first (a–c) and seventh (d–f, early-learning) visit. The magenta dashed and the blue solid curves in (d) correspond to a normal and GMM fit with three components, respectively. The kurtosis (*κ*) values and the total number of trajectories (*N*) analyzed are indicated next to the respective panels in (a–f). The dataset analyzed here was collected at 100 Hz. For the present study, we focused on the data during the pre-feeding phases only. (g–l) Histograms of active exploratory velocity at low (g–i: 15 lx) and high (j–l: 300 lx) illumination level in three different species of hawkmoth—nocturnal *Deilephila elpenor* (g,j), diurnal *Macroglossum stellatarum* (h,k), and crepuscular *Manduca sexta* (i,l). Colors of the fits are same as in (a–f). The kurtosis (*κ*) values and the total number of hawkmoths (*N*) analyzed are indicated. (m–o) Box and whisker plots showing the RMS active exploratory velocity for all three species of hawkmoths analyzed at different illumination levels. The one-tailed p-values were computed using the Mann-Whitney-Wilcoxon test. The datasets analyzed here were collected at 100 Hz with one trial per moth.

**Extended Data Fig. 6.**
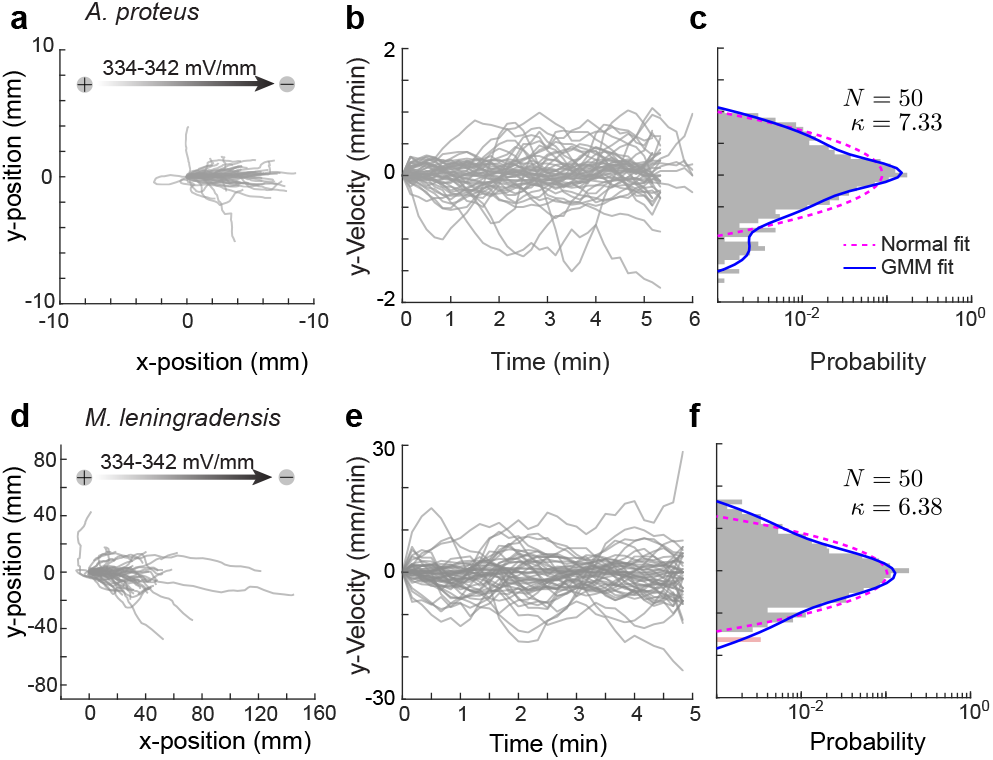
Reanalysis of the directed movement of *Amoeba proteus* and *Metamoeba leningradensis* in response to an electric field (galvaontaxis).^22^. Migration trajectories (*N* = 50) of *A. proteus* (a), temporal traces of velocities (b) in the transverse direction of the applied electric field derived from the migration data, and the corresponding velocity histogram (c) with kurtosis (*κ*) value. The magenta dashed and the blue solid curves in (c) correspond to a normal and GMM fit with three components, respectively. (d-f) Galvanotaxis response in *M. leningradensis*. The panels are same as in (a-c). Image data for both datasets were collected at 0.1 Hz.

**Extended Data Fig. 7.**
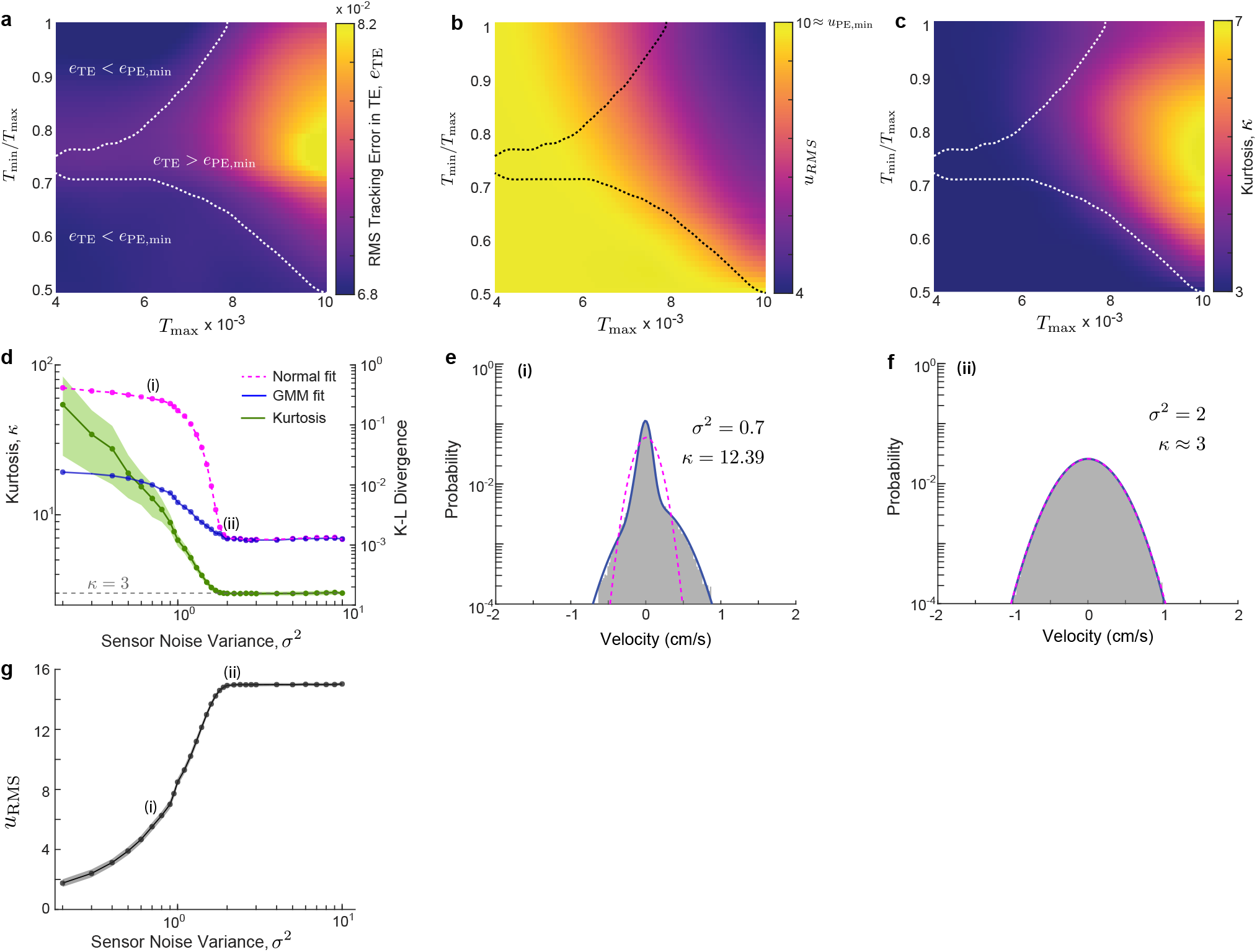
Effect of thresholds, *T*_min_ and *T*_max_ in Triggered Excitation strategy. (a–c) Heatmaps showing mean RMS tracking error in Triggered Excitation (*e*_TE_, a), mean RMS control effort (*u*_RMS_, b) and mean kurtosis (*κ*, c) of the resultant velocity distributions from 100 independent simulations at critical excitation level corresponds to minimum RMS tracking error in Persistent Excitation (*e*_PE,min_ in Fig. 5l) as thresholds *T*_min_ and *T*_max_ were varied. The dashed line in (a–c) shows the phase transition based on the difference between the tracking error in Triggered and Persistent Excitation. The region inside the line corresponds to parameter space where the tracking error in Persistent Excitation is less than Triggered excitation, whereas outside the region Triggered excitation performs better. (d) Variation of kurtosis, *κ* (green, left *y*-axis), and Kullback–Leibler (K-L) divergence (right *y*-axis) of normal distribution (magenta dashed) and Gaussian mixture model (blue solid) fit to the velocity distribution with sensor noise variance, *σ*^2^. (e,f) Velocity histograms with kurtosis values are shown for sensor noise variance, *σ*^2^ = 0.70 and 2, respectively, as indicated by (i) and (ii) in (d). (g) Variation of RMS control effort (*u*_RMS_) with sensor noise variance, *σ*^2^. The shaded regions in (d,g) denote the respective standard deviations (*n* = 25 independent simulations per *σ*^2^)

**Extended Data Table 1:**
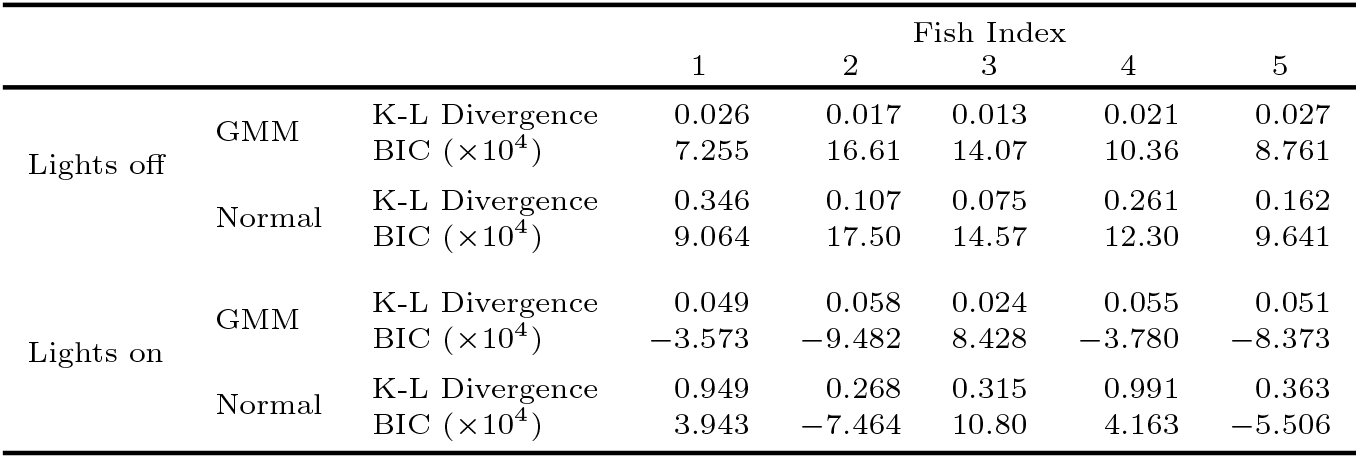
Comparison of Kullback-Leibler (K-L) divergence and Bayesian information criterion (BIC) for the three-component Gaussian mixture model (GMM) and normal fits to the velocity data of *Eigenmannia virescens* during lights-off and lights-on trials. GMM with lower values of K-L divergence and BIC is favored over normal.

**Extended Data Table 2:**
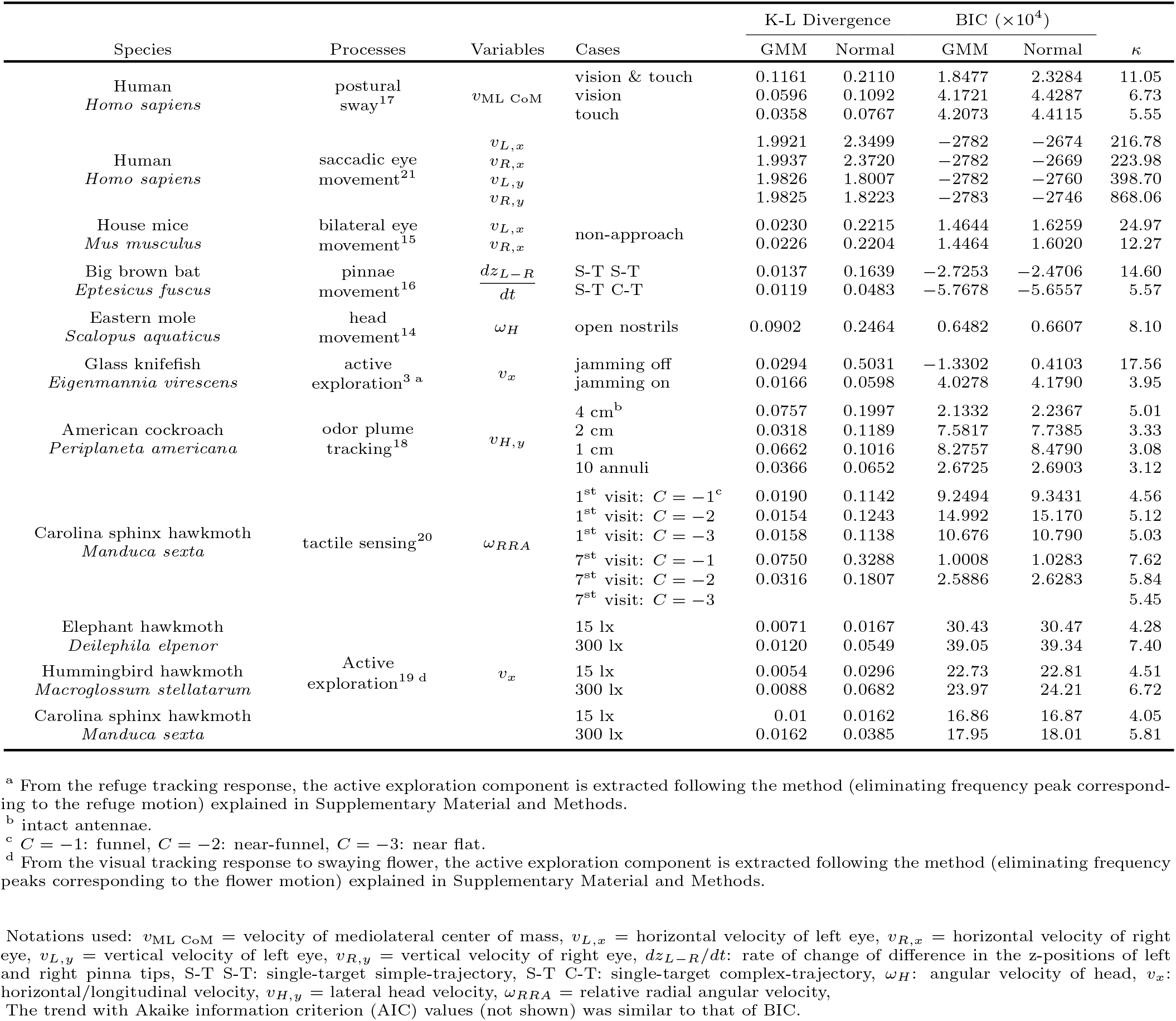
Summary of the reanalysis of the published data across nine different species^3^,^14–21^ showing Kullback–Leibler (K-L) divergence and Bayesian information criterion (BIC) for the normal and the three-component Gaussian mixture model (GMM) fits to the velocity data and the kurtosis value (*κ*).

**Extended Data Table 3:**
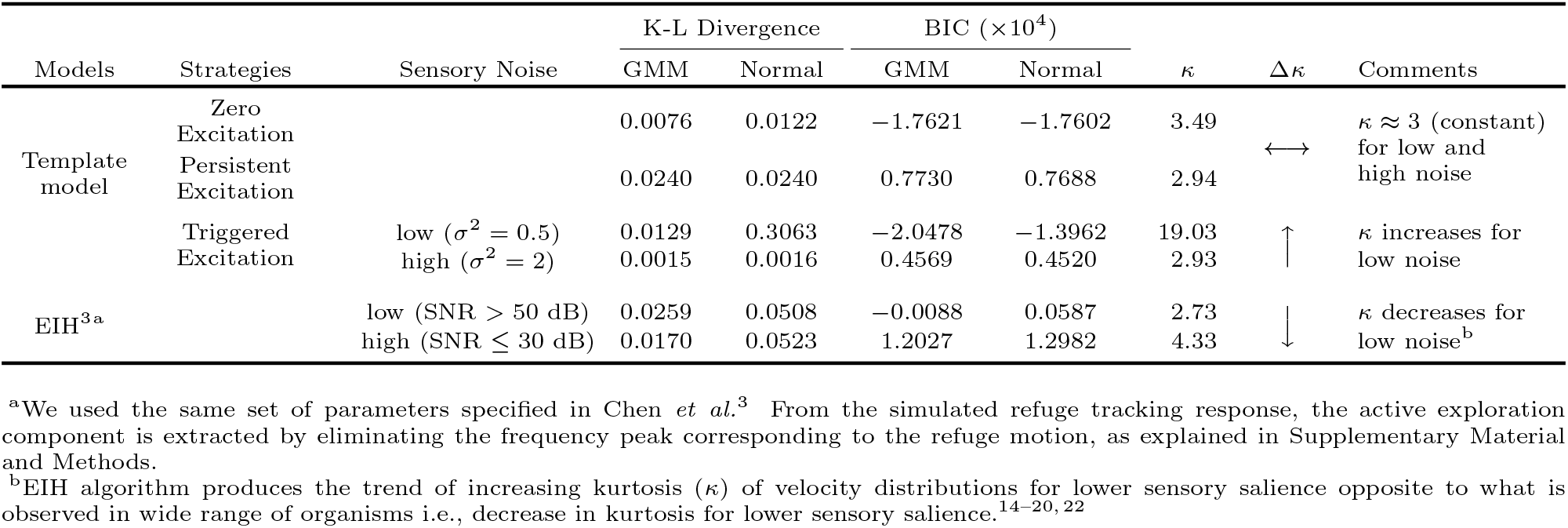
Summary of comparison between different strategies of template model and Ergodic Information Harvesting (EIH) model showing Kullback-Leibler (K-L) divergence and Bayesian information criterion (BIC) for the normal and the three-component Gaussian mixture model (GMM) fits to the simulated velocity data and the kurtosis value (*κ*).

**Supplementary Fig. 1.**
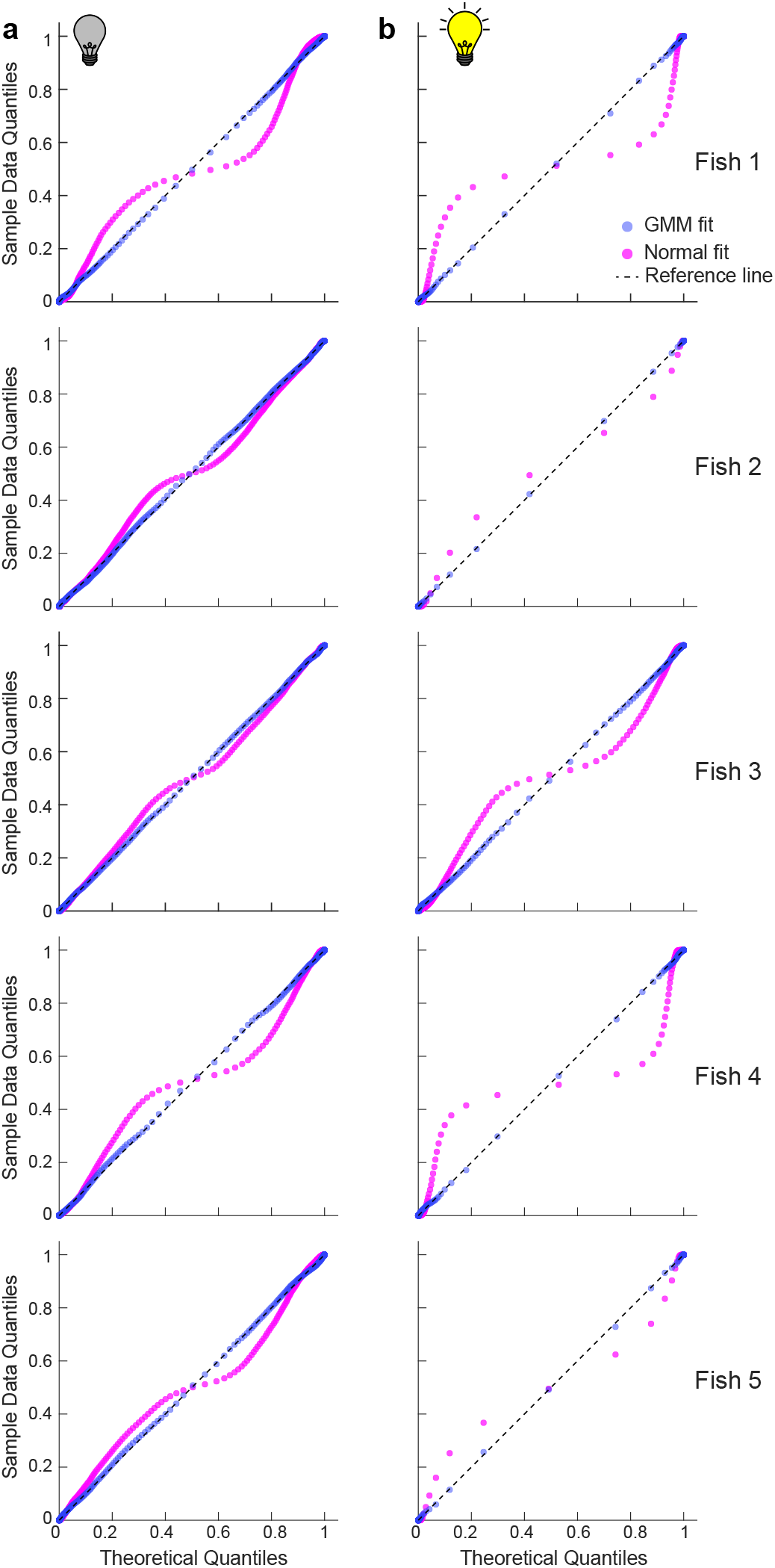
Q-Q plots of velocity traces are consistent with mode-switching. (a, b) Q-Q plots comparing the velocity data for each fish (*N* = 5) from lights-off (a) and lights-on trials (b) with theoretical quantiles from a normal fit (magenta) and GMM fit (blue), respectively. Lesser deviation from the reference line (black dashed) for GMM fits with three components than for the normal, suggested better fitting of the former. The Q-Q plots also showed that the lights-off trial data were closer to normal distribution fits than were those of lights-on trial data.

**Supplementary Fig. 2.**
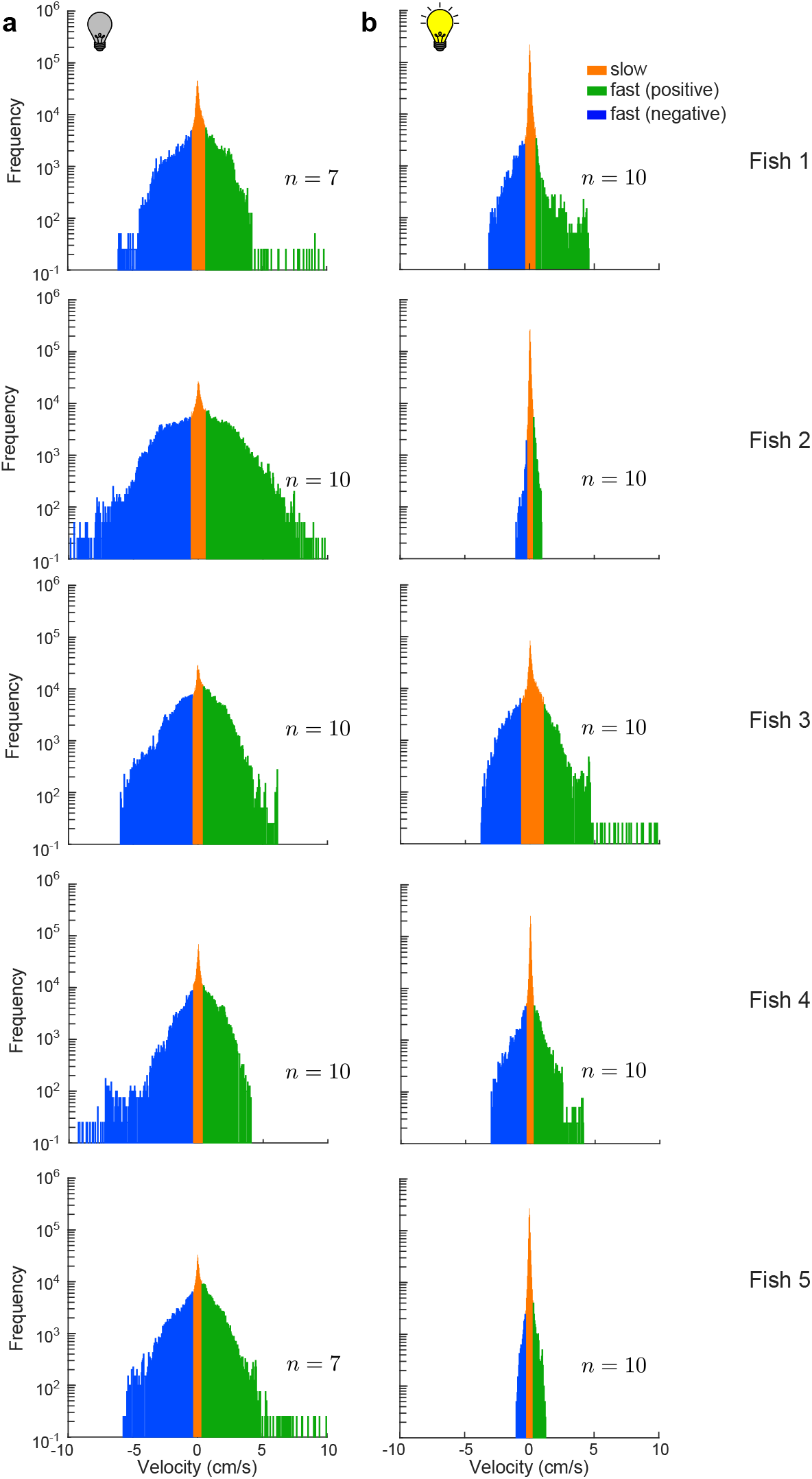
Clustering of velocity data into different behavioral modes. (a,b) Velocity histograms of lights-off (a) and lights-on (b) trials with three clusters: slow (orange), fast positive (green) and fast negative (blue). The clustering was based on GMM model with inflection point based algorithm (see Methods for details).

**Supplementary Fig. 3.**
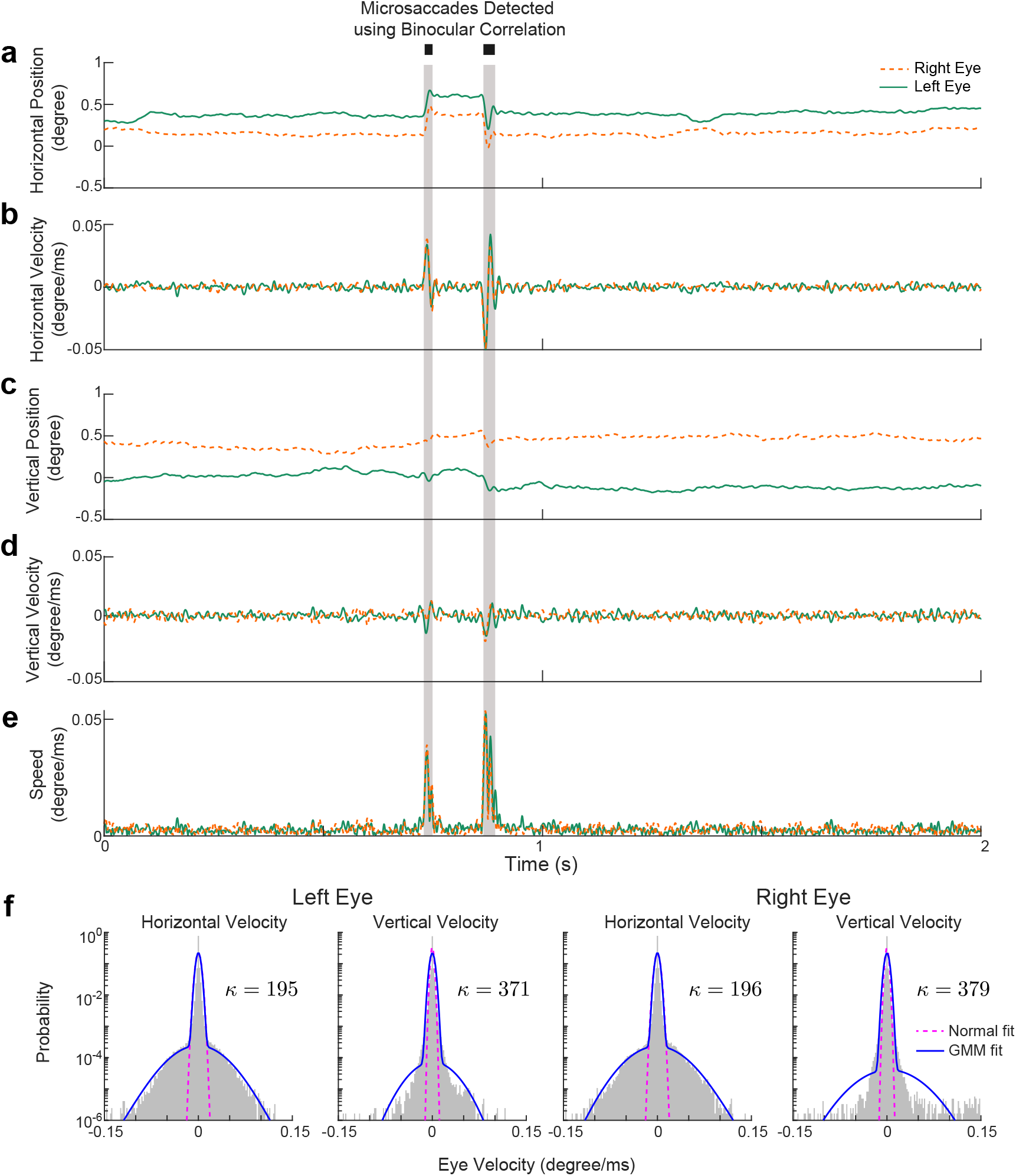
Reanalysis of the gaze fixation in humans, *Homo sapiens*.^21^. (a–e) Representative temporal traces of horizontal eye position (a) and velocity (b), vertical eye position (c) and velocity (d), and eye speed (e) of right (orange dashed) and left (green) eye. The gray sections denote the detected microsaccades using the binoculer correlation (BC) method proposed in Hauperich *et al.*^21^ (f) Velocity histograms of horizontal and vertical velocities of left and right eye with respective kurtosis (*κ*) values. The magenta dashed and the blue solid curves correspond to a normal and GMM fit with three components, respectively. The dataset was collected from five human adults at 1000 Hz.

**Supplementary Fig. 4.**
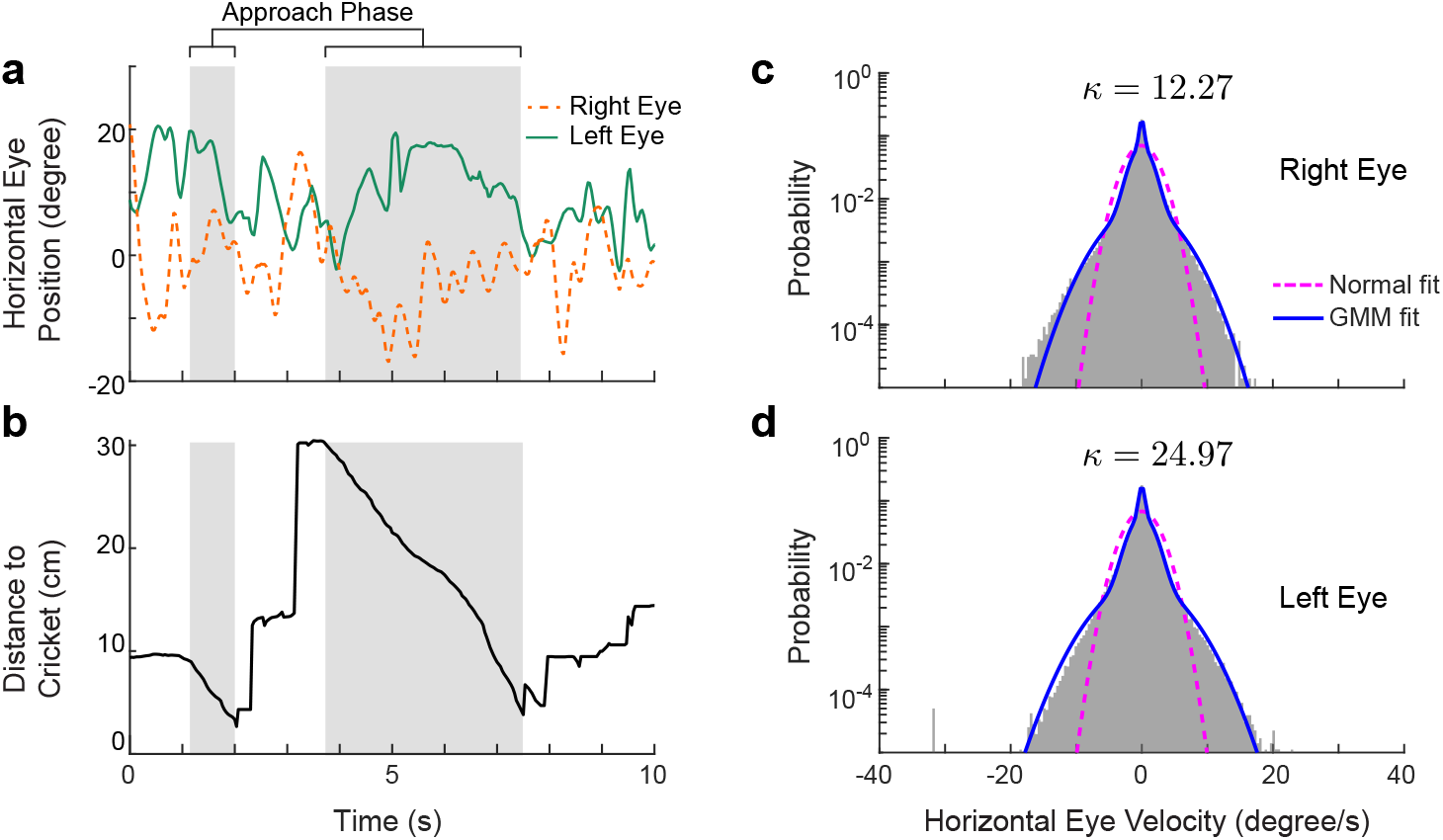
Reanalysis of the bilateral eye movements in house mice, *Mus musculus* during prey capture.^15^. (a) Representative temporal traces of horizontal eye positions of right (orange dashed) and left (green) eye. (b) Representative plot of distance between the mouse and the cricket over time. The grey boxes denote the approach phases during which the mouse head was oriented towards the cricket (the azimuthal angle was with in *±*45*^o^*) and was moving at a speed over 1 cm*/*s. For the present study, we focused on the data during the non-approach phases (*∼* 95% of the total data). (c,d) Histograms of horizontal eye velocities of right (c) and left (d) eye during the non-approach phases with respective kurtosis (*κ*) values. The magenta dashed and the blue solid curves correspond to a normal and GMM fit with three components, respectively. The dataset has 105 trajectories, collected from seven mice at 30 Hz.

**Supplementary Fig. 5.**
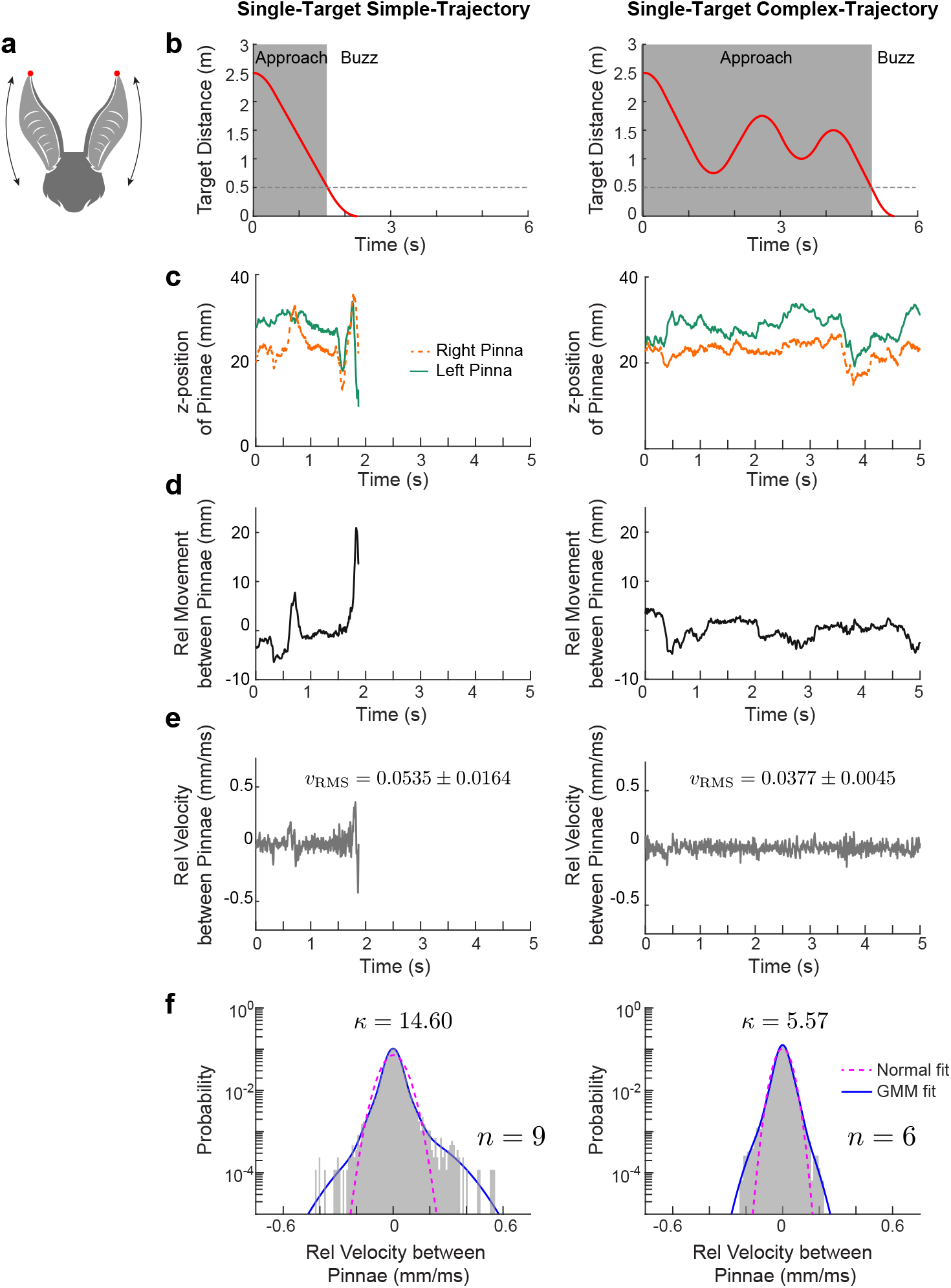
Reanalysis of the tip of the pinnae movement in big brown bats, *Eptesicus fuscus*.^16^. (a) Cartoon showing the “head-waggle” behavior where bat changes the relative z-positions of the tip of its pinnae (red dots). (b) The simple (left) and the complex trajectory (right) of the tethered target (mealworm) over time. The gray box denotes the “approach” phase during which the target distance was greater than 0.5 m (dashed line). The rest we considered as the “buzz” phase. A sharp drop in the pulse duration and the pulse interval marked the beginning of the buzz phase and we found that the distance threshold 0.5 m roughly correlates to the onset of the buzz phase. For the present study, we focused on the data during the approach phase only (ignoring the last 200 frames). (c-f) Representative temporal traces of change in the z-positions (c) of the tip of the pinnae (orange dashed: right pinna, green: left pinna), relative movement (difference in z-positions) between pinnae (d), relative velocity between pinnae (e), and the histograms of relative velocity (f) with respective kurtosis (*κ*) values and the total number of trials (*n*) analyzed for the cases of the simple (left) and the complex trajectories (right). Mean *±* SD of the RMS values of the relative velocity (*v*_RMS_) are shown next to plots in (e). The one-tailed p-value = 0.004 for *v*_RMS_, was computed using the Mann-Whitney-Wilcoxon test. The magenta dashed and the blue solid curves in (f) correspond to a normal and GMM fit with three components, respectively. The dataset has trajectories collected from three bats at 500 Hz.

**Supplementary Fig. 6.**
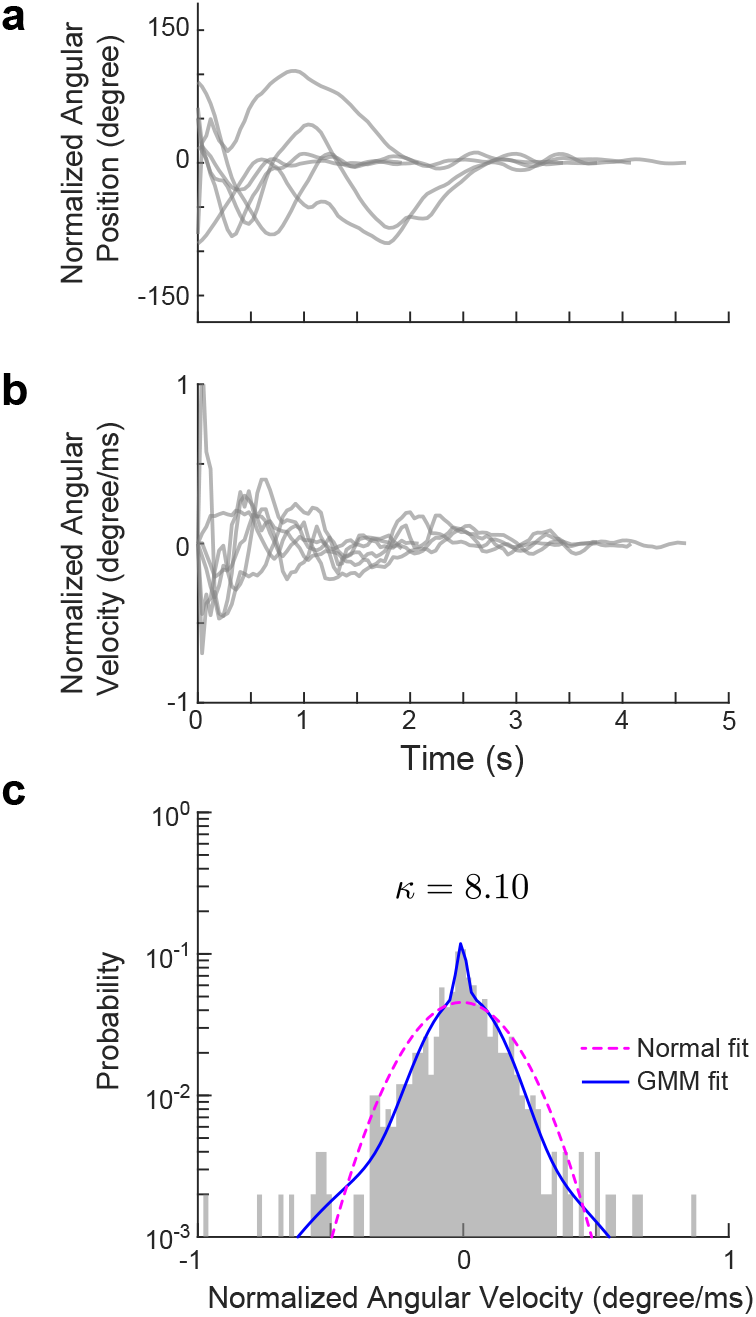
Reanalysis of the olfactory response in Eastern moles, *Scalopus aquaticus*.^14^. (a) Temporal traces (*n* = 6) of angular position of moles normalized such that the destination direction was 0*^◦^*. (b) Traces of angular velocity (assuming the data was sampled at 25 Hz). (c) Histograms of angular velocities with the kurtosis (*κ*) value. The magenta dashed and the blue solid curves correspond to a normal and GMM fit with three components, respectively.

## Footnotes

a Active sensing includes not only the sorts of ancillary movements described here, but also the emission of energetically costly signals such as electric fields by weakly electric fishes^42^ and echolocation calls in dolphins, birds, and bats.^43–46^ The emission and reception of these electrical and acoustic signals are also affected by movements of the individual, which has been a focus of our work in electric fishes.^2, 5, 8^

## References

[1] Gibson, J. J. Observations on active touch. Psychol Rev 69, 477–491 (1962).

[2] Biswas, D. et al. Closed-loop control of active sensing movements regulates sensory slip. Curr Biol 28, 4029–4036 (2018).

[3] Chen, C., Murphey, T. D. & MacIver, M. A. Tuning movement for sensing in an uncertain world. eLife 9, e52371 (2020).

[4] Soatto, S. Actionable information in vision. In Machine learning for computer vision, 17–48 (Springer, 2013).

[5] Stamper, S. A., Roth, E., Cowan, N. J. & Fortune, E. S. Active sensing via movement shapes spatiotemporal patterns of sensory feedback. J Exp Biol 215, 1567–1574 (2012).

[6] Cowan, N. J. & Fortune, E. S. The critical role of locomotion mechanics in decoding sensory systems. J Neurosci 27, 1123–1128 (2007).

[7] Rose, G. J. & Canfield, J. G. Longitudinal tracking responses of the weakly electric fish, *Sternopygus*. J Comp Physiol A 171, 791–798 (1993).

[8] Uyanik, I., Stamper, S. A., Cowan, N. J. & Fortune, E. S. Sensory cues modulate smooth pursuit and active sensing movements. Front Behav Neurosci 13, 59 (2019).

[9] Sefati, S. et al. Mutually opposing forces during locomotion can eliminate the tradeoff between maneuverability and stability. Proc Natl Acad Sci U.S.A. 110, 18798–18803 (2013).

[10] Golovin, D. & Krause, A. Adaptive submodularity: Theory and applications in active learning and stochastic optimization. J Artif Intell Res 42, 427–486 (2011).

[11] Cooper, G. F. The computational complexity of probabilistic inference using Bayesian belief networks. Artif Intell 42, 393–405 (1990).

[12] Blondel, V. D. & Tsitsiklis, J. N. A survey of computational complexity results in systems and control. Automatica 36, 1249–1274 (2000).

[13] Narendra, K. S. & Annaswamy, A. M. Persistent excitation in adaptive systems. Int J Control 45, 127–160 (1987).

[14] Catania, K. C. Stereo and serial sniffing guide navigation to an odour source in a mammal. Nat Commun 4, 1–8 (2013).

[15] Michaiel, A. M., Abe, E. T. & Niell, C. M. Dynamics of gaze control during prey capture in freely moving mice. eLife 9, e57458 (2020).

[16] Wohlgemuth, M. J., Kothari, N. B. & Moss, C. F. Action enhances acoustic cues for 3-D target localization by echolocating bats. PLoS Biol 14, e1002544 (2016).

[17] Kiemel, T., Oie, K. S. & Jeka, J. J. Multisensory fusion and the stochastic structure of postural sway. Biol Cybern 87, 262–277 (2002).

[18] Lockey, J. K. & Willis, M. A. One antenna, two antennae, big antennae, small: total antennae length, not bilateral symmetry, predicts odor-tracking performance in the American cockroach *Periplaneta americana*. J Exp Biol 218, 2156–2165 (2015).

[19] Stöckl, A. L., Kihlström, K., Chandler, S. & Sponberg, S. Comparative system identification of flower tracking performance in three hawkmoth species reveals adaptations for dim light vision. Philos Trans R Soc Lond B Biol Sci 372, 20160078 (2017).

[20] Deora, T., Ahmed, M. A., Daniel, T. L. & Brunton, B. W. Tactile active sensing in an insect plant pollinator. J Exp Biol 224, jeb239442 (2021).

[21] Hauperich, A.-K., Young, L. K. & Smithson, H. E. What makes a microsaccade? A review of 70 years of research prompts a new detection method. J Eye Mov Res 12 (2019).

[22] De la Fuente, I. M. et al. Evidence of conditioned behavior in amoebae. Nat Commun 10, 1–12 (2019).

[23] Sutton, E. E., Demir, A., Stamper, S. A., Fortune, E. S. & Cowan, N. J. Dynamic modulation of visual and electrosensory gains for locomotor control. J R Soc Interface 13, 20160057 (2016).

[24] Kunapareddy, A. & Cowan, N. J. Recovering observability via active sensing. In Proc Amer Control Conf, 2821–2826 (IEEE, Milwaukee, WI, USA, 2018).

[25] Sontag, E. D., Biswas, D. & Cowan, N. J. An observability result related to active sensing (2022). URL https://arxiv.org/abs/2210.03848.

[26] Fabre, M. et al. Large postural sways prevent foot tactile information from fading: Neurophysiological evidence. Cereb Cortex Comm 2 (2020).

[27] Arkley, K., Grant, R. A., Mitchinson, B. & Prescott, T. J. Strategy change in vibrissal active sensing during rat locomotion. Curr Biol 24, 1507–1512 (2014).

[28] Matthews, M. & Sponberg, S. Hawkmoth flight in the unsteady wakes of flowers. J Exp Biol 221, jeb179259 (2018).

[29] Tritico, H. M. & Cotel, A. J. The effects of turbulent eddies on the stability and critical swimming speed of creek chub *Semotilus atromaculatus*. J Exp Biol 213, 2284–2293 (2010).

[30] Harris, C. M. & Wolpert, D. M. Signal-dependent noise determines motor planning. Nature 394, 780–784 (1998).

[31] Nelson, M., Xu, Z. & Payne, J. Characterization and modeling of p-type electrosensory afferent responses to amplitude modulations in a wave-type electric fish. J Comp Physiol A 181, 532–544 (1997).

[32] Lee, J. et al. Templates and anchors for antenna-based wall following in cockroaches and robots. IEEE Trans Robot 24, 130–143 (2008).

[33] Jun, J. J., Longtin, A. & Maler, L. Active sensing associated with spatial learning reveals memory-based attention in an electric fish. J Neurophysiol 115, 2577–2592 (2016).

[34] Clarke, S. E., Naud, R., Longtin, A. & Maler, L. Speed-invariant encoding of looming object distance requires power law spike rate adaptation. Proc Natl Acad Sci U.S.A. 110, 13624–13629 (2013).

[35] Clarke, S. E., Longtin, A. & Maler, L. A neural code for looming and receding motion is distributed over a population of electrosensory on and off contrast cells. Journal of Neuroscience 34, 5583–5594 (2014).

[36] Takeda, K. et al. Incoherent feedforward control governs adaptation of activated ras in a eukaryotic chemotaxis pathway. Science signaling 5, ra2–ra2 (2012).

[37] Biswas, D., Devreotes, P. N. & Iglesias, P. A. Three-dimensional stochastic simulation of chemoattractant-mediated excitability in cells. PLoS computational biology 17, e1008803 (2021).

[38] Bertsekas, D. Dynamic programming and optimal control: Volume I, vol. 1 (Athena scientific, 2012).

[39] Mobbs, D., Trimmer, P. C., Blumstein, D. T. & Dayan, P. Foraging for foundations in decision neuroscience: insights from ethology. Nat Rev Neurosci 19, 419–427 (2018).

[40] Kaelbling, L. P., Littman, M. L. & Moore, A. W. Reinforcement learning: A survey. J Artif Intell Res 4, 237–285 (1996).

[41] Zweifel, N. O. & Hartmann, M. J. Defining “active sensing” through an analysis of sensing energetics: homeoactive and alloactive sensing. J Neurophys 124, 40–48 (2020).

[42] Bullock, T. H. Electroreception. Annu Rev Neurosci 5, 121–170 (1982).

[43] Brinkløv, S., Elemans, C. P. & Ratcliffe, J. M. Oilbirds produce echolocation signals beyond their best hearing range and adjust signal design to natural light conditions. R Soc Open Sci 4, 170255 (2017).

[44] Nelson, M. E. & MacIver, M. A. Sensory acquisition in active sensing systems. J Comp Physiol A 192, 573–586 (2006).

[45] Snyder, J. B., Nelson, M. E., Burdick, J. W. & MacIver, M. A. Omnidirectional sensory and motor volumes in electric fish. PLoS Biol 5, e301 (2007).

[46] Ghose, K. & Moss, C. F. Steering by hearing: a bat’s acoustic gaze is linked to its flight motor output by a delayed, adaptive linear law. J Neurosci 26, 1704–1710 (2006).

[47] Berg, H. C. & Brown, D. A. Chemotaxis in *Escherichia coli* analysed by three-dimensional tracking. Nature 239, 500–504 (1972).

[48] Benda, J., Longtin, A. & Maler, L. A synchronization-desynchronization code for natural communication signals. Neuron 52, 347–358 (2006).

[49] Metzen, M. G., Hofmann, V. & Chacron, M. J. Neural synchrony gives rise to amplitude-and duration-invariant encoding consistent with perception of natural communication stimuli. Front Neurosci 14, 79 (2020).

[50] Hofmann, V. & Chacron, M. J. Population coding and correlated variability in electrosensory pathways. Front Integr Neurosci 12, 56 (2018).

[51] Grewe, J., Kruscha, A., Lindner, B. & Benda, J. Synchronous spikes are necessary but not sufficient for a synchrony code in populations of spiking neurons. Proc Natl Acad Sci U.S.A. 114, E1977–E1985 (2017).

[52] Fetcho, J. R., Higashijima, S.-i. & McLean, D. L. Zebrafish and motor control over the last decade. Brain Res Rev 57, 86–93 (2008).

[53] Tabor, K. M. et al. Direct activation of the Mauthner cell by electric field pulses drives ultrarapid escape responses. J Neurophysiol 112, 834–844 (2014).

[54] Orchard, G. et al. Hfirst: A temporal approach to object recognition. IEEE Trans Pattern Anal Mach Intell 37, 2028–2040 (2015).

[55] Hitschfeld, É. M., Stamper, S. A., Vonderschen, K., Fortune, E. S. & Chacron, M. J. Effects of restraint and immobilization on electrosensory behaviors of weakly electric fish. ILAR journal 50, 361–372 (2009).

[56] Roth, E., Zhuang, K., Stamper, S. A., Fortune, E. S. & Cowan, N. J. Stimulus predictability mediates a switch in locomotor smooth pursuit performance for *Eigenmannia virescens*. J Exp Biol 214, 1170–1180 (2011).

[57] Vágvolgyi, B. P. General tracker. https://github.com/vagvolgyi/general_tracker (2021).

[58] Ratcliffe, J. M., Elemans, C. P., Jakobsen, L. & Surlykke, A. How the bat got its buzz. Biol Lett 9, 20121031 (2013).

